# Molecular architecture of the developing mouse brain

**DOI:** 10.1101/2020.07.02.184051

**Authors:** Gioele La Manno, Kimberly Siletti, Alessandro Furlan, Daniel Gyllborg, Elin Vinsland, Christoffer Mattsson Langseth, Irina Khven, Anna Johnsson, Mats Nilsson, Peter Lönnerberg, Sten Linnarsson

**Author notes:** Equal contribution (Gioele La Manno, Kimberly Siletti). Correspondence to: G.L. ( / @GioeleLaManno / gioelelamanno.com) S.L. ( / @slinnarsson / linnarssonlab.org).

## Abstract

The mammalian brain develops through a complex interplay of spatial cues generated by diffusible morphogens, cell-cell interactions, and intrinsic genetic programs that result in the generation of likely more than a thousand distinct cell types. Therefore, a complete understanding of mammalian brain development requires systematic mapping of cell states covering the entire relevant spatiotemporal range. Here we report a comprehensive single-cell transcriptome atlas of mouse brain development spanning from gastrulation to birth. We identified almost a thousand distinct cellular states, including the initial emergence of the neuroepithelium, a rich set of region-specific secondary organizers and a complete developmental program for the functional elements of the brain and its enclosing membranes. We used the atlas to directly test the hypothesis that human glioblastoma reflects a return to a developmental cell state. In agreement, most aneuploid tumor cells matched embryonic rather than adult types, while karyotypically normal cells predominantly matched adult immune cell types.

---

**The brain emerges** from the primitive ectoderm as a sheet of neuroepithelial cells which folds into the neural tube during neurulation^1^. The developing nervous system is unique for the length of the developmental window, the extent of the interplay between different anatomical regions and lineages, and the diversity of cell types generated. Therefore, the ability of single-cell RNA-seq to disentangle the molecular heterogeneity of a complex cell pool has been particularly useful to study nervous system development^2–10^. Recent studies have shed light on the developing telencephalon^5,11^, the hippocampus^9,12,13^, the developing ventral midbrain^14–16^, the developing spinal cord and cerebellum^17,18^, and the hypothalamic arcuate nucleus and diencephalon^19,20^. Single-cell RNA-seq has elucidated the differences between embryonic, postnatal and adult neural progenitors^9,21,22^, and compared normal glial progenitors with their malignant counterparts^23,24^.

To map mouse brain development in detail, we collected embryonic brain tissue from 43 pregnant CD-1 mice, sampling each day from E7 to E18 (Extended Data Figure 1a-b, Table S1). We prepared 105 samples by droplet-based single-cell RNA sequencing. After removing low-quality cells and doublets (Methods), 96 samples remained with a mean of 5766 transcripts (unique molecular identifiers, UMIs) and 1 934 genes detected per cell (Extended Data Figure 1c-f). The total cellular RNA content dropped as a function of embryonic age in all lineages, and then increased again in the most mature neurons (Extended Data Figure 1e-f and Extended Data Figure 2c-d). Erythrocytes segregated into primitive and definitive erythropoiesis and showed exceptionally high UMI counts but low gene counts (Extended Data Figure 2a). A transcriptional signature of cell-cycle activation (Methods) showed a clear bimodal pattern where 32% of all cells were cycling (>1% cell-cycle gene UMIs; Figure 1 and Extended Data Figure 2b). Immature cells from young embryos were generally proliferating, whereas more mature cells from older embryos had stopped dividing. Postmitotic cells expressed a larger proportion of nascent (unspliced) RNA relative to mature (spliced) RNA (Extended Data Figure 2e), likely due to the induction of lineage-specific genes when cells began their maturation programs.

**Figure 1.**
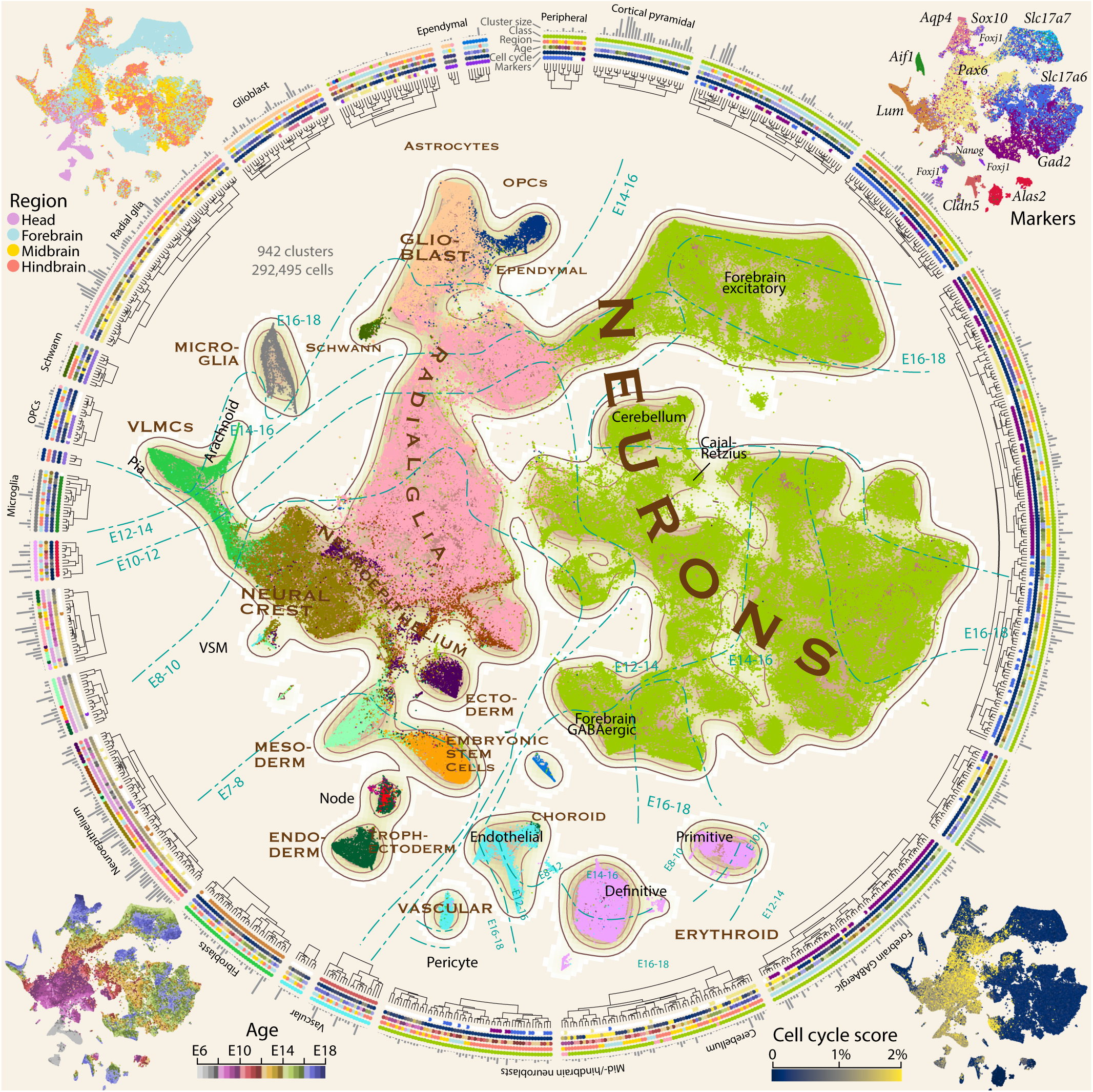
Atlas of the developing mouse nervous system. All 292,495 high-quality cells are shown using a tSNE embedding (center) colored by major classes as indicated in brown capital lettering. Dashed curves show approximate regions of gestational pseudoage intervals. The four corner insets show the dissected region (upper left), the expression of key marker genes (upper right), the cell-cycle score (lower right) and the pseudoage (lower left; defined as the average gestational age of a cell and its 25 nearest neighbors). The surrounding radial dendrogram shows all 942 clusters. The dendrogram was cut for clarity into 25 major categories of cells as indicated by gaps. Grey bars indicate cluster size (number of cells), and four colored tracks indicate class (colored as the central tSNE), region, pseudoage, cell cycle and marker gene expression (each colored as the corner tSNEs).

A tSNE embedding of the dataset (Figure 1) revealed that cells were organized primarily by gestational age, branching to show the major lineages originating from the neuroepithelium. A large connected component — representing the neural tube and its derivatives — was surrounded by disconnected islets of microglia, erythrocytes, and vascular cells, none of which are derived from the neuroepithelium. From E7 the embryo split into clusters corresponding to the three germ layers, then formed the early neural tube. By E8-10, the neural crest had formed and from E10 to E16 differentiated into the VLMC (vascular and lep-tomeningeal cells^25^, also known as brain fibroblast-like cells^26^) lineage which proceeded to form the two layers of the meninges, pia and arachnoid^27^. The early neural tube then matured into proliferating radial glia, the precursors of all neural cells. Two large cohorts of neurons branched from the neural tube, one comprising forebrain excitatory neurons, and one containing all other neuronal types. The clear separation of forebrain excitatory neurons agrees with our previous findings in the adult mouse brain^22^, indicating the strong transcriptional and functional specialization of those cell types. After about E14, radial glia gradually lost proliferative capacity and switched to a glioblast state that eventually gave rise to astrocytes, ependymal cells and oligodendrocyte precursor cells. The orderly progression of successive waves of cell types reveal the coordinated temporal development of the mammalian brain (Extended Data Figure 2f).

**Focusing first on** the late gastrulation and early neurulation stages, we extracted E7-E8 cells from our dataset (Figure 2a-b) and divided the population into three parts (Figure 2b-c): primitive streak-stage cells (*Pou5f1* and *T*, ref. 28); non-neural lineage-restricted progenitor cells (*Foxa2* and *Krt8*, refs. 29,30); and early neuroepithelial cells (*Pax6* and *Hes3*, refs. 31,32). Aligning our dataset to a previously described single-cell atlas of gastrulation (Figure 2d-f, ref. 33) confirmed that our data captured a continuous set of cell states from gastrulation and neurulation (Figure 2e) but revealed a greater heterogeneity of states than previously reported (Figure 2f).

**Figure 2.**
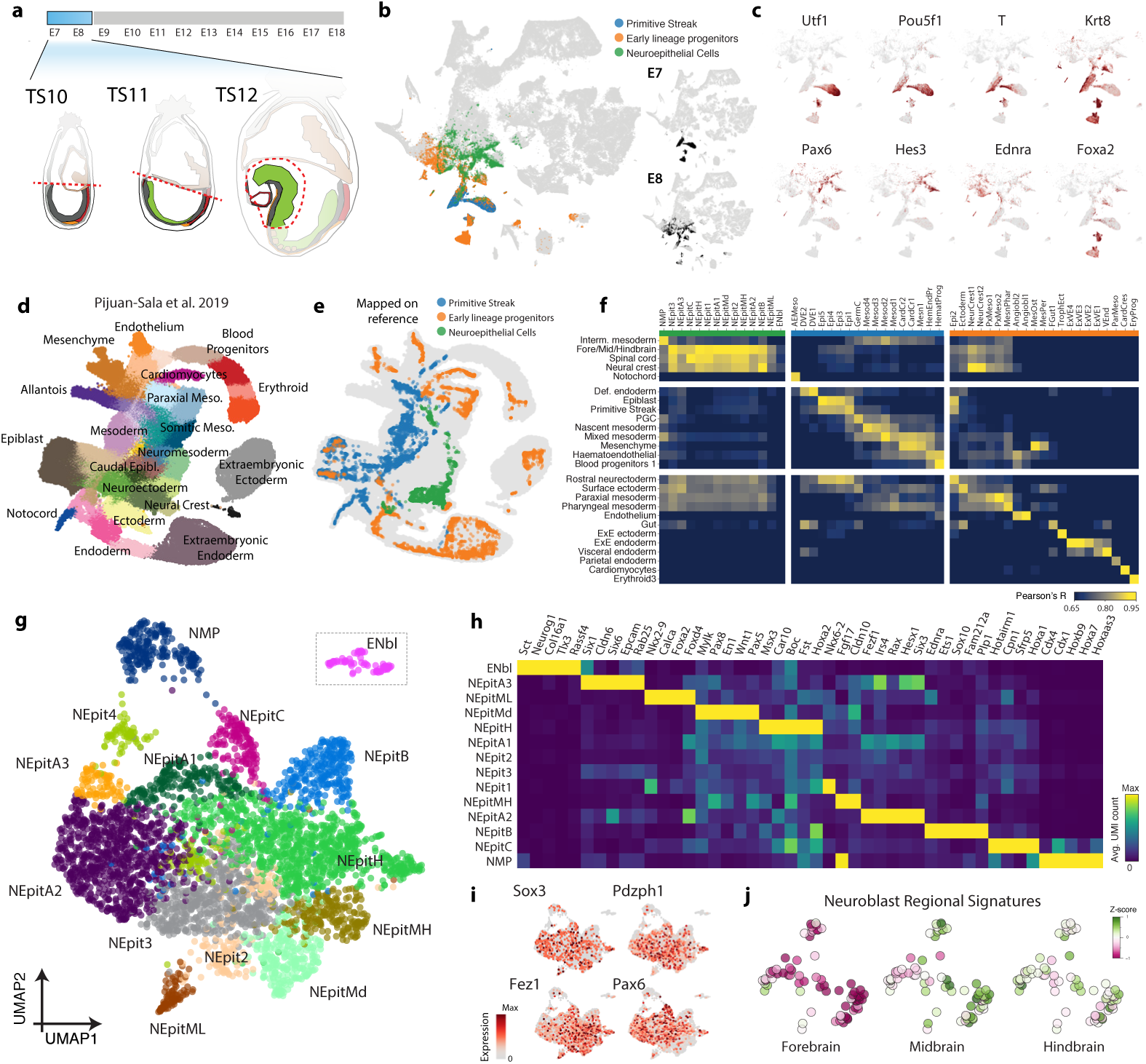
Emergence of cell type heterogeneity from gastrulation to neurulation. **a**, Scheme of tissue sampling for early neurulation timepoints (E7-E8). **b**, The three subsets of cells sampled at early time points (E7-E8) overlaid on the global tSNE embedding of Figure 1. **c**, Expression of lineage-specific marker genes overlaid to the global tSNE as in (b). **d**, tSNE map of the mouse gastrulation single-cell atlas from Pijuan-Sala et al. **e**, Gene expression data from primitive streak cells, non-neural lineage-specific progenitors and neuroepithelial cells projected on the embedding describing mouse gastrulation33. **f**, Pairwise correlation between neurulation-stage clusters of this paper with those of Pijuan-Sala et al. **g**, UMAP embedding of the neuroepithelial cells obtained from timepoints E7-8. Cells are colored by clusters. Dashed boxes indicate that the ENbl cluster was repositioned as it was located further in the embedding space. **h**, A heatmap displaying gene expression of enriched genes in each of the neuroepithelial populations. **i**, Expression of broad neuroepithelial markers overlaid on the UMAP in (g). **j**, Patterning-related gene signature scores computed for the cells in the earliest neuroblast cluster (ENbl). Signatures consisted of genes enriched from all the other neuroblast cluster that were sampled from different regions of the brain.

Among neural cells we identified fourteen populations: twelve neuroepithelial clusters expressing *Fez1, Sox3, Pax6*, and *Pdzph1*^34^, one neuromesodermal progenitor cluster expressing *Cdx4, Cdx1* and a set of caudal *Hox* genes^35,36^ and a population of early neuroblasts expressing *Nhlh1* and *Nhlh2*^37^ (Figure 2g-i). The twelve neuroepithelial populations comparably expressed a set of stem-cell genes but differentially expressed genes linked to spatial patterning (Figure 2h, refs. 38,39). We recognized anterior populations including forebrain, anterior neural ridge and eye field progenitors (NEpitA1-3), a population expressing midbrain, hindbrain and spinal cord pattering factors (NEpitMd, NEpitMH, NEpitH and NEpitC) but also populations with a stark mediolateral patterned signature corresponding to the progenitors located at the midline and the neural fold (NEpitML and NEpitB). Finally, the population of early neuroblasts showed heterogeneous gene expression and likely represented a mixture of cells. Comparing these cells with later neuroblasts, revealed an enrichment of a midbrain and hindbrain signature with telencephalic-specific genes being less represented, suggesting that the first neuroblasts appear posteriorly (Figure 2j; Methods).

**Next, we examined** the neurulation and morphogenesis stages, E9-E11. Differently from gastrulation, where signals for the establishment of the neural plate anteroposterior axis are provided extrinsically, during neurulation specific neural progenitors, called secondary organizers, orchestrate the morphogenesis of the brain. We examined the distribution of genes encoding morphogens including SHH, WNTs, BMPs, FGFs, Neuregulins and R-spondins and found them enriched in a subset of E9-E11 radial glia-like cells (Figure 3a-b; Extended Data Figure 5a-b). We found four ventralizing organizers expressing *Shh*, four dorsalizing organizers expressing *Wnts* and *Bmps*, three *Fgf*-expressing organizers that characterized cells at the boundaries between regions, and two *Neuregulin*-expressing antihem organizers that appeared later in development (Figure 3c-d). The localization of these organizers clusters was corroborated by querying the Allen Developmental Mouse Brain ISH Atlas using Voxhunt (Extended Data Figure 5c; Methods; ref. 40).

**Figure 3.**
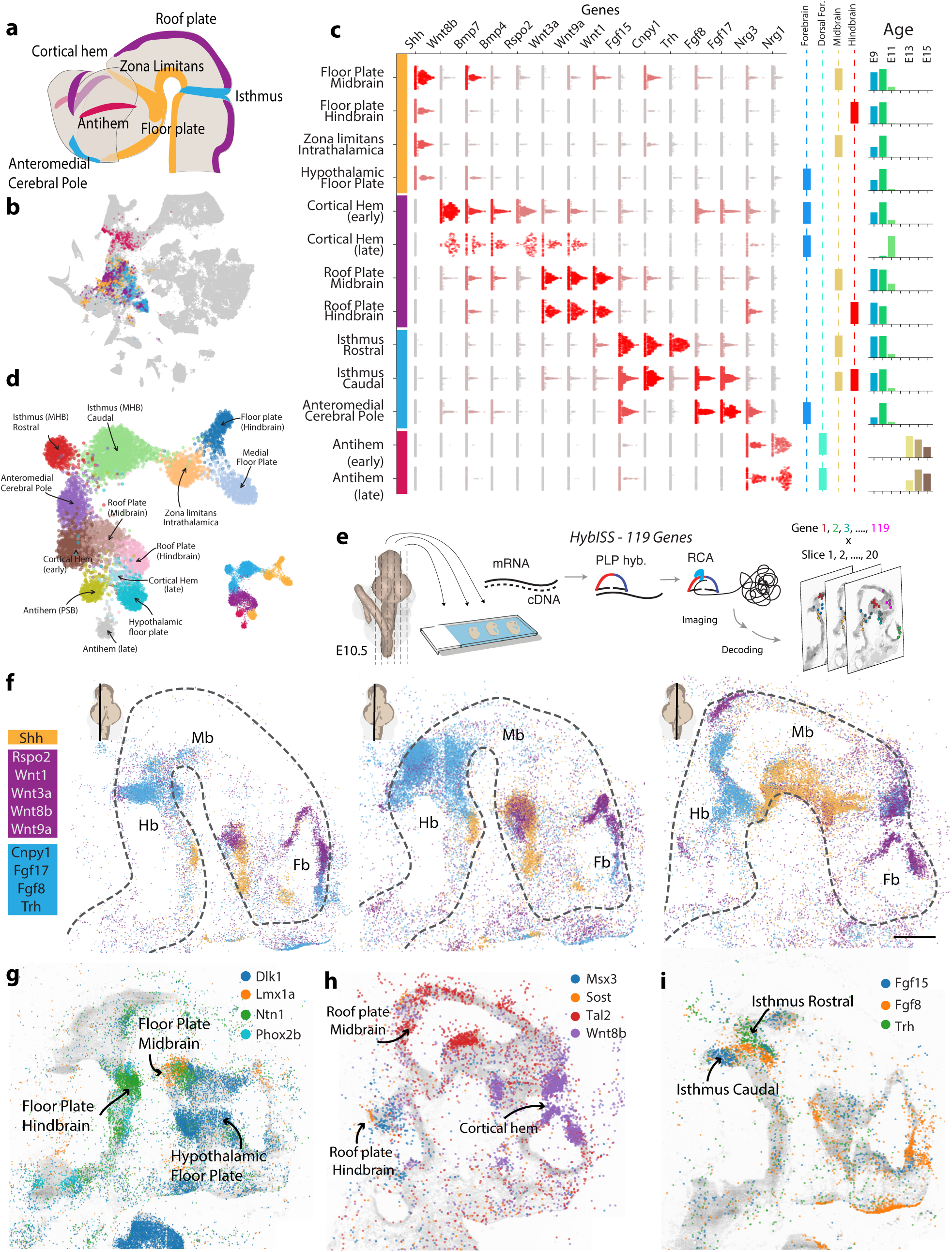
Characterization of molecular identity and signal-receptor repertoire of neural tube organizers. **a**, A schematic representation of the neural tube marking the localization of secondary organizers. Colors group early organizers by ventralizing (yellow), dorsalizing (violet) and boundary-inducing (cyan) activity. Late organizers are in fuchsia. **b**, The cells belonging to organizers-related clusters are overlaid on the global tSNE of Figure 1. **c**, Level of expression of receptors and ligands genes described in the literature as defining organizers. Beeswarm plots are showing distribution of expression (log-transformed) for each cluster, each dot is a cell, cells are colored by the average expression in that cluster. **d**, UMAP embedding of radial-glia like cells that we identified as related to specific organizers. The miniature displays the same embedding colored by organizer type, as defined in (a). **e**, A scheme of the in situ sequencing approach used to localize the expression of the 119 genes listed in (c) and Extended Figure 5b-c. **f**, The expression of a selection of organizer-specific genes E10.5 embryo, as detected by in-situ sequencing. Dots represent the localized transcripts of different genes and are colored by organizer type. Miniatures indicate the approximate mediolateral position of the section. Dashed lines demarcate the neural tube. Fb, forebrain; Mb, midbrain; Hb, hindbrain. Scale bar 480 μm. **g**, Expression of selected genes that demarcate the distention between different kind of floor plate h, dorsal i, and boundary organizers.

Because organizer cells were identified by clustering, their transcriptomes were distinct from those of other cells and not defined only by the small number of genes described in previous literature. A large number of transcription factors and secreted ligands and surface receptors were expressed specifically in subtypes of organizers (Extended Data Figure 5d-e). For example, *Wnt7b* and *Rspo2* distinguished dorsalizing organizers in the cortical hem, whereas *Wnt3* and *Fzd10* were specific to roof plate organizers. *Ntf3* and *Sost* were expressed by mesencephalic and rhombencephalic roof plate cells, respectively. All dorsalizing organizers expressed *Wnt9a, Wnt3a, Resp1, Resp3* and *Bmp6*.

## Box 1 Resources

The raw sequence data is deposited in the sequence read archive under accession PRJNA637987.

The companion wiki at http://mousebrain.org provides the following resources:

- An interactive version of Figure 1.
- Expression matrices.
- Metadata

The analysis software developed for this paper is available at https://github.com/linnarsson-lab, in repositories named *cytograph, punchcards* and *auto-annotation-md*.

Though defined by *Shh* expression, each of the ventralizing organizers were further distinguished by region-specific expression of secreted molecules. The hypothalamic floor plate was the only ventralizing organizer that expressed *Dlk1*, a Notch ligand, and was distinguished by the absence of *Spon1* and *Ntn1*. Furthermore, the Zona limitans intrathalamica (ZLI) and the mesencephalic floor plate shared expression of *Wnt5a, Wnt5b* and *Lrtm1* but were distinguished by *Pappa* and *Dkk2* expression, respectively. These results establish a rich repertoire of secondary organizers secreting combinations of morphogens that may act region-specifically to induce locally distinct neuronal fates.

To directly validate these secondary organizers, we performed in-situ sequencing^41^ of 119 genes (Figure 3c and Extended Data Figure 5d-e) in an E10.5 mouse brain (Figure 3e-i). The spatial distributions of *Shh, Wnt*, and *Fgf* isoforms clearly revealed the secondary organizers (Figure 3f). Moreover, the subtypes of floor, roof, and boundary organizers identified by scRNA-seq were revealed as spatial domains of the more general organizers (Figure 3g-i), defining a more complex system of spatial regulation of brain development than previously recognized. To summarize the complexity of the 119 spatial patterns of expression we clustered the in-situ sequencing data defining 40 embryonic territories characterized by a unique repertoire of signaling molecules and patterning factors (Extended Data Figure 6a-b). Overall, the fact that subtypes of organizers were both transcriptionally distinct, spatially segregated, and expressed distinct sets of ligands and receptors, shows that progenitors born in even closely adjacent spatial domains are subject to distinct environmental cues.

**A distinct clade** of the cluster dendrogram comprised late glia (Figure 1; clusters 751 and above) and related progenitors (Figure 4a and Extended Data Figure 7a-f, Methods). Focusing on these cells revealed two major groups of radial glia: those that expressed neurogenic markers, like *Neurog2* or *Dlx2*, and those that expressed glial markers like *Tnc* or *Egfr* (Figure 4b-c). We refer to the latter as glioblasts, given their likely commitment to gliogenesis. More than 6 000 genes were significantly enriched in one of the two groups (Figure 4d, Extended Data Figure 7c-d). Over 600 of these genes were cell-cycle related, 89% of which were enriched in neurogenic radial glia. *Olig1, Olig2, Aqp4, Stat3*, and glutamate and GABA receptors were enriched in glioblasts. To further distinguish progenitors related to astrocytes or oligodendrocyte precursor cells (OPCs), we compared with genes enriched in these cell types in the adolescent mouse (Extended Data Figure 8j).

**Figure 4.**
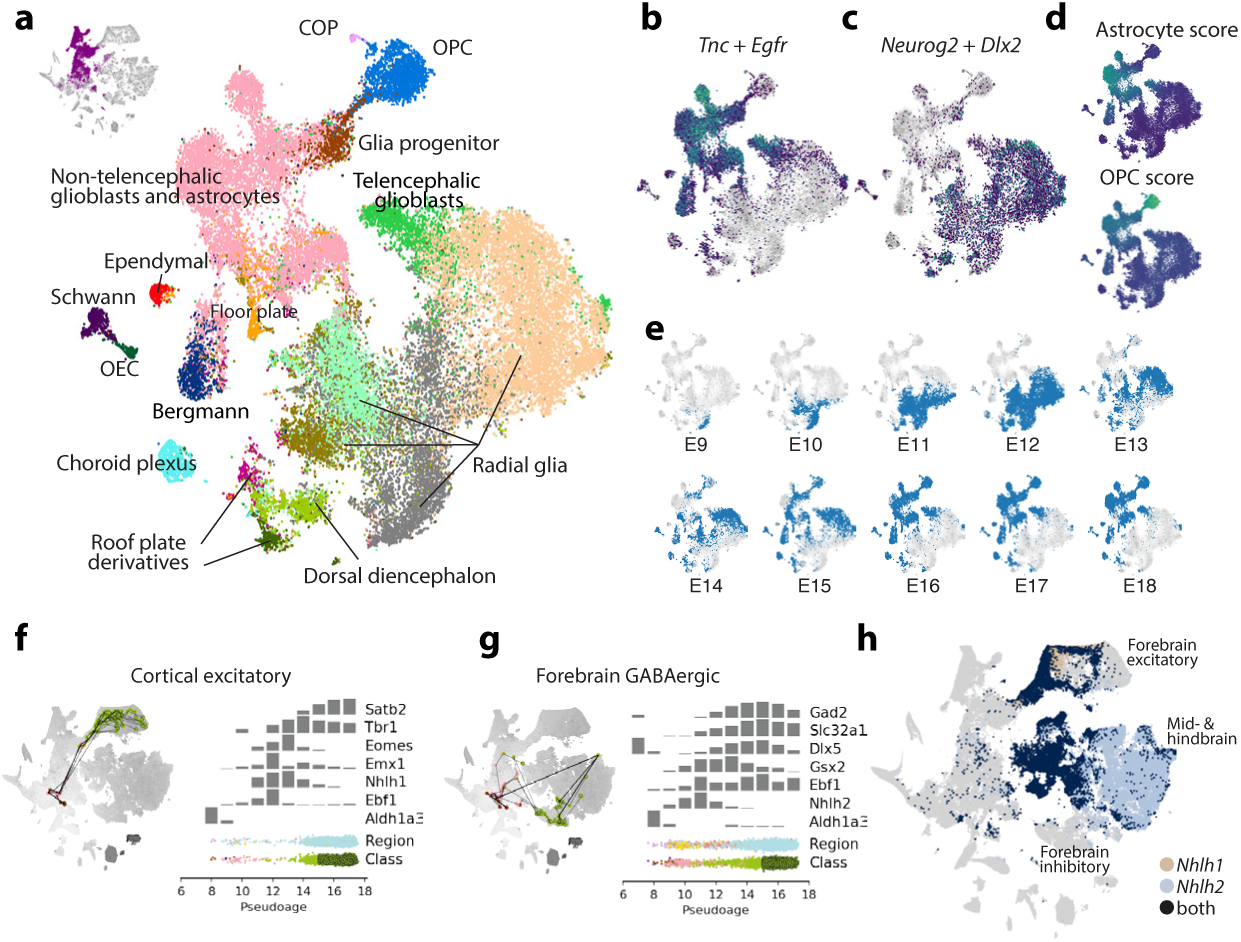
Glioblasts and neuroblasts. **a**, Clusters in the last clade of the dendrogram, representing glioblasts and related cell types, were analyzed separately (inset shows the cells that were included). A new dendrogram was calculated and cut to result in 18 clusters, labeled and indicated by colors. **b**, Cells on the tSNE colored according to the sum of pro-neural *Neurog2* and *Dlx2* expression. **c**, Cells on the t-SNE colored according to the sum of *Tnc* and *Egfr* expression. **d**, Gestational age of cells on the tSNE. **e**, Pseudo-lineages of cortical excitatory cells. **f**, Pseudolineage of forebrain GABAergic cells. **g**, Expression of neuroblast markers, transcription factors *Nhlh1* and *Nhlh2*.

We previously defined seven region-specific astrocyte types of the adolescent mouse brain^22^, dominated by abundant telencephalic and non-telencephalic subtypes. Consistent with this observation, non-telencephalic glioblasts and oligodendrocyte precursors of mixed forebrain, midbrain, and hindbrain origin were distinct from a cluster of predominantly telencephalic glioblasts biased towards astrocytes (Extended Data Figure 7e). In contrast, neurogenic radial glia split into separate clusters based on region. These observations suggest a loss of regionalization particularly in the oligodendrocyte lineage, but also among astrocytes. To investigate further, we sampled from our dataset six groups of cells - early glia, neurogenic radial glia, glioblasts, OPCs, neuroblasts, and neurons – and trained a classifier to identify a cell’s region of origin using transcription-factor expression (Extended Data Figure 7g-i). The classifier was trained on each group and tested on the five remaining groups. Although the results suggest that region-defining genes shift throughout development, the classifier still performed relatively well when trained on glioblasts and OPCs, demonstrating that some region-defining genes were still present at low levels (Extended Data Figure 7j).

The transition from neurogenic to gliogenic occurred between E12 and E16 (Figure 4e and Extended Data Figure 7e). The earliest gliogenic cell to appear was a cluster unique in its co-expression of *Egfr* and the Notch ligands *Dll1, Dll3*, and *Dll4* (Extended Data Figure 7k-l), This cluster, which was closely related to OPCs and might represent a pre-OPC state, expressed high levels of *Olig1* and *Olig2*, as well as *Crlf1*, a soluble member of the *Cntfr* pathway implicated in gliogenesis^42^. The appearance of astrocytes at approximately E15 was revealed by increased levels of *Gfap, Agt*, and *Aqp4* (Extended Data Figure 7k).

**To better resolve** lineage-specific versus shared gene expression programs, we inferred a pseudolineage tree for the whole dataset (Extended Data Figure 8). We arbitrarily designated a single cell in the early neuroepithelium as the root, and computed shortest paths (i.e., geodesics on the manifold) to every other cell on the radius nearest neighbor graph (RNN). The set of all shortest paths from the root forms a pseudolineage tree, due to the convergence of paths towards the root (Extended Data Figure 8a-b). Plotting individual pseudolineage trajectories to a few selected single cells (Extended Data Figure 8c-d) on the atlas embedding confirmed that pseudolineages agreed with expectations. For example, pia and arachnoid originated from the cranial neural crest; astrocytes and OPCs were generated from distinct but closely related radial glia; and cortical excitatory neurons and forebrain GABAergic neurons were derived from distinct radial glia precursors.

The pseudolineage tree provides a tool to investigate gene expression along putative lineage trajectories leading to specific end states. To do this, we isolated all the trajectories that ended in a specific cluster (or set of clusters), and projected all the single cells in those trajectories onto the embryonal pseudoage (Figure 4f-h and Extended Data Figure 8e-i). We then calculated average gene expression in pseudoage bins. To validate this approach, we first examined the subtree terminating in cortical excitatory neurons (Figure 4f) whose development is well understood. As expected, cells along the trajectory predominantly were located in the forebrain. Furthermore, a set of known regulators of cortical excitatory neurogenesis was expressed in an ordered progression from E11 to E18: *Emx1, Eomes, Tbr1* and *Satb2*^11^. RNA in situ hybridization (Allen Brain Atlas) confirmed the timing of expression of these key transcription factors (Extended Data Figure 8k).

Expanding the analysis to additional glial and neuronal lineages showed the expected lineage markers (Figure 4g and Extended Data Figure 8e-i). We noticed that nearly all neuronal — but none of the glial — lineages passed through an early neuroblast state expressing all or most of *Nhlh1, Nhlh2, Ebf1, Ebf2* and *Ebf3*, extending the HPF analysis above with lineage-specificity. The expression of *Nhlh1* and *Nhlh2*, especially in combination, (Figure 4g, Extended Data Figure 8j) identified a pan-neuronal state just at or after cell cycle exit (Figure 1 and Extended Data Figure 2b), which coincided with the set of clusters we had manually annotated as neuroblasts. Notably, however, the forebrain GABAergic lineage appeared to lack *Nhlh1* expression, although low levels of *Nhlh2* were detected.

**Human brain cancer** is a devastating disease, with a median survival of less than two years for glioblastoma. Recent work using single-cell RNA-sequencing to determine the cellular composition of glioblastoma has suggested that these tumors reflect a reversion to an embryonic cell state resembling neural progenitors, OPCs and/or immature astrocytes^24,43,44^. To unambiguously test this hypothesis, we compared human brain tumor data — comprising five glioblastoma samples and one anaplastic astrocytoma^45^ — with a reference cell type catalog constructed by merging the developmental cell atlas of this paper with our previous adolescent brain cell atlas^22^ (Methods and Extended Data Figure 9).

Each of the tumor samples matched a unique set of reference cell types, but in each case aneuploid cells predominantly matched embryonic cell types (mainly radial glia and neuroblasts) or adult astrocytes (Figure 5). In contrast, normal (euploid) cells matched adult immune cells, oligodendrocytes and vascular cells. Euploid vascular cells invariably matched embryonic vascular cells, not adult, likely indicating ongoing angiogenesis. The tumors differed in composition, proliferative status and the nature of the immune response. For example, only the anaplastic astrocytoma showed B cell and T cell response, and the most highly proliferative glioblastoma (SF11215) was almost devoid of immune cells. However, every tumor contained proliferating progenitors resembling embryonic radial glia or OPCs. With one exception (a neural match in tumor SF11159), all aneuploid clusters that matched adult cell types were astrocyte, OPC or neuroblast-like; all of which are cell types that are phenotypically close to embryonic progenitor cells. Thus, an unbiased comparison with embryonic and adult cell types confirms the essentially fetal cellular nature of human glioblastoma.

**Figure 5.**
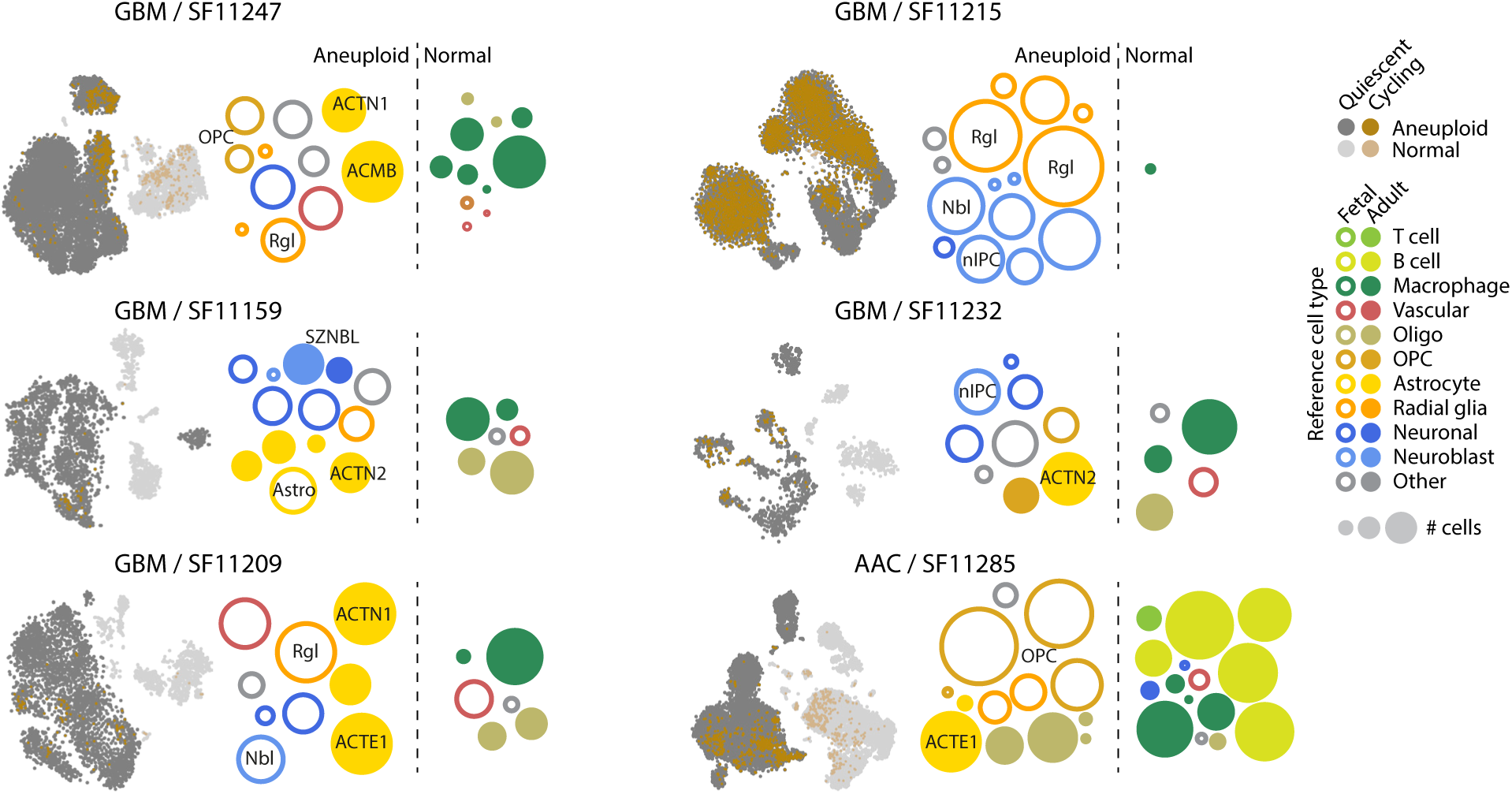
Developmental nature of human glioblastoma. Each subpanel shows a tSNE plot of tumor cells, indicating aneuploid (dark grey) and cycling (brown) cells, and bubbles indicating the best match for each tumor cluster to a reference cell type (942 types from this paper and 265 types from Zeisel et al. 2018), colored by major class. Open circles are developmental reference types; closed circles are adolescent reference types; bubbles are positioned arbitrarily. Immune cell types were manually called, as they were not present in the reference cell type compendium. GBM, glioblastoma multiforme. AAC, anaplastic astrocytoma.

**Our atlas is** a step toward dissecting the principles of mammalian nervous-system development and provides an overview of the prenatal brain’s transcriptomic diversity. The wealth of information on time-, lineage-, and region-specific gene expression provides powerful tools for genetic targeting, and for understanding genes involved in neurodevelopmental disorders and human brain cancer.

## Acknowledgements

We thank Hannah Hochgerner and Amit Zeisel (Technion, Israel) for valuable advice, Patrik Ernfors (Karolinska Institute) for supporting the work of A.F., and the National Genomics Infrastructure for sequencing services. This work was supported by grants from the Knut and Alice Wallenberg Foundation (2015.0041 to S.L. and 2018.0172 to S.L. and M.N.), the Erling Persson Family Foundation (HDCA, to S.L. and M.N.), the Swedish Foundation for Strategic Research (RIF 15-0057, SB16-0065) to S.L., Hjärnfonden (PS2018-0012) to D.G., the Chan Zuckerberg Initiative DAF, an advised fund of Silicon Valley Community Foundation (2018-191929) to M.N, and the Swiss National Science Foundation (CRSK-3_190495) to G.L.

## Author contributions

S.L. supervised the project. S.L. and G.L. conceived the study design. G.L., K.S. and S.L. analyzed, annotated and interpreted the single-cell data and wrote the manuscript. G.L. and A.F. performed the single-cell experiments. A.F. performed the embryo dissociations. A.J. prepared sequencing libraries. P.L. and S.L. built the companion website. E.V., D.G. and C.M.L. performed in situ sequencing experiments. K.S., I.K., G.L. and C.M.L. analyzed and interpreted the in situ sequencing data. M.N. supervised the *in situ* sequencing experiments. All the authors critically reviewed the manuscript and approved the final version.

## Code and data availability

Source code is available at https://github.com/linnarsson-lab. RNA-seq data is available at the Sequence Read Archive (SRA; https://www.ncbi.nlm.nih.gov/sra) under accession PRJNA637987. HybISS data is available at http://mousebrain.org/downloads.

## Methods

### Animals

CD-1 mice were obtained from Charles River (Germany) and mated overnight. The morning that plugs were detected was considered E0.5. Mice were housed with regular dark/light cycle and fed standard diet and water ad libitum. All animal procedures were approved by the Stockholm ethics committee (N68/14; Stockholms djurförsöksetiska nämnd) and followed Directive 2010/63/EU of the European Parliament and of the Council, the Swedish Animal Welfare Act (Djurskyddslagen: SFS 1988:534), the Swedish Animal Welfare Ordinance (Djurskyddsförordningen: SFS 1988:539) and the provisions regarding the use of animals for scientific purposes: DFS 2004:15 and SJVFS 2012:26.

### Dissections

161 embryos between E7 and E18.5 were collected from 36 pregnant mothers. Embryos were extracted from the uterine horn and dissected in phosphate buffered saline (PBS) on ice within one hour. Dissections were performed with microsurgical tools under a fluorescent stereomicroscope. The number of embryos dissected was adapted to accommodate changes in size and cell numbers in the developing brain (see Table S1).

E7 and E8 embryos were cut into two parts to isolate the rostro-ventral neural tube from dorsal tissue including the trophectoderm, which was discarded. The developing spinal cord was identified by corresponding somite pairs and also discarded. From E9 and E10 embryos, we isolated the entire cephalic part, including prospective forebrain, midbrain, and hindbrain. From E11 onwards, only developing brain tissue was dissected and collected; surrounding head tissue was discarded. Meninges was removed whenever possible. E9 to E11 brains were divided into forebrain, midbrain and hindbrain. Forebrain and midbrain were separated at the diencephalon; midbrain and hindbrain at the isthmus. E12 to E15 forebrains were further divided into dorsal (including the hemispheres) and ventral (including the subcortical territories, caudoputamen and the ventral striatum) parts. E16 to E18 forebrains were divided into dorsal, ventral, and thalamic parts.

### Cell dissociation

Dissected tissue was processed as previously described, with minor modifications^9,22^. Tissue was transferred to a 2 ml Eppendorf tube containing preheated digestion solution (37°C). Before E11 Trypsin was used for dissociation; after E11 the digestion solution contained 300 µl TrypLE Express (Life Technologies; cat. no. 12605-010), 2 000 µl papain solution (Worthington Biochemical; cat. no. LK003178; 25 U/ml in cutting solution), 100 µl DNase I (Worthington Biochemical; cat. no. LK003172; 1 mM in cutting solution) and 100 µl cutting solution. Enzymatic incubation time was adjusted based on developmental stage. The tissue was kept in a 37°C water bath and gently pipetted three times every 10 minutes. After 30 min at 37°C, another 100 µl of TrypLE solution was added, followed by 10 more minutes at 37°C and pipetting. After approximately 45–50 min, dissociation was complete, and cells became visible at the bottom of the Eppendorf tube. The solution was filtered using a 20-µm cell strainer (Falcon cat. no. 352340) and collected in a 15-ml plastic tube. The digestion solution was diluted with 2.4 ml of cutting solution and 0.6 ml of Neurobasal-A medium (Gibco; cat. no. 10888) and centrifuged at 100g for 4 min at 4°C. The supernatant was removed and the pellet resuspended in 0.5 ml cutting solution and 0.5 ml Neurobasal-A medium. Neurobasal-A medium was supplemented with l-glutamine (Gibco; cat. no. 25030-123), B27 (Gibco; cat. no. 17504-044) and penicillin plus streptomycin (Sigma 50x; cat. no. P4458). The cell suspension was carefully transferred with a Pasteur pipette onto an Optiprep gradient. For the gradient, 85 µl of Optiprep Density Solution (Sigma; cat. no. D1556) was mixed with 457.5 µl cutting solution and 457.5 µl Neurobasal medium including supplements. The gradient was subsequently centrifuged at 80 g for 10 min at 4°C. The supernatant was removed until only 100–200 µl remained. DNase I (10 µl) was added to prevent aggregation. The single-cell density was evaluated. Then cells were filtered with a 20 µm strainer (Partec CellTrics). Cells were pelleted, resuspended in cutting solution with DNaseI, and loaded into 10X Genomics Chromium v1 chips.

### Single-cell RNA sequencing

Droplet-based single-cell RNA sequencing was performed using the 10x Genomics Chromium Single Cell Kit v1. Single-cell suspensions concentrated at 500-700 cells/ml were mixed with master mix and nuclease free water according to the Chromium manual, targeting 3500 cells per reaction. 12 PCR cycles were used for cDNA synthesis. Each library was sequenced using an Illumina HiSeq instrument, one sample per lane, with one 98 bp read located near the 3′ end of the mRNA. Illumina runs were demultiplexed and aligned to the genome (mm10-3.0.0), and BAM files were obtained from the 10X Genomics cellranger pipeline (version 3.0.2). Molecule counts were attributed to spliced and unspliced transcripts by running velocyto^46^ (version 0.17.11) with standard parameters, resulting in one loom file (http://loompy.org) per sample.

### In-situ sequencing by HybISS

HybISS^41^ was performed as published at protocols.io (http://dx.doi.org/10.17504/protocols.io.xy4fpyw). Padlock probes, bridge probes and detection oligo sequences used are listed in Supplementary Table 3. Imaging was performed with a Leica DMi8 epifluorescence microscope equipped with LED light source (Lumencor® SPECTRA X), sCMOS camera (Leica DFC9000GTC), and 20× objective (HC PL APO, 0.80). ROIs were imaged with 10% overlap and 24 z-stack planes with 0.5 μm spacing and then maximum projected in LASX software and then tiles exported as raw TIFF files.

### Computational Analysis

#### Pooling samples and preprocessing for analysis

Loom files from each 10x sample were aggregated by time point and brain region, the latter dictated by our dissection strategy. In particular we pooled: Cephalic Neural Tube E7-8, Forebrain E9-11, Midbrain E9-11, Hindbrain E9-11, Forebrain Dorsal E12-15, Forebrain Ventral E12-15, Midbrain E12-15, Hindbrain E12-15, Forebrain Dorsal E16-18, Forebrain Ventrothalamic E16-18, Forebrain Ventrolateral E16-18, Midbrain E16-18, and Hindbrain E16-18. Cells with fewer than 2 000 UMIs were excluded from pooling. Genes expressed in fewer than ten cells or greater than 60% of cells were excluded from analysis.

### Cytograph 2.0

The analysis pipeline is available as an update to our Cytograph package, Cytograph 2.0 (https://github.com/linnarsson-lab/cytograph-dev). Cytograph is under continuous development, and the results in this paper were generated at commit 2e8eb79a5f83b-d9dda15503d9b2476ebdd8fa0b3. We described Cytograph v1.0 elsewhere^22^, including gene enrichment (overexpression), trinariation and clustering. We detail below only the key improvements: changes to dimensionality reduction and graph construction; auto-annotation; and punchcards.

In brief, we used Hierarchical Poisson factorization (HPF^47^) to decompose the expression matrix into 96 non-negative and modular components, while reducing noise and preserving most of the structural information in the original matrix. We then computed a radius nearest neighbor graph (RNN) using the information radius (also called the Jensen-Shannon divergence, a measure of the Shannon information difference) to link cells with near-identical gene-expression states. Finally, we clustered the RNN graph using modularity (Louvain) to define distinct sets of cells representing cell types or states along differentiation trajectories. We thus represent the high-dimensional gene expression manifold simultaneously as an RNN graph (facilitating analysis of continuous processes such as differentiation), a set of clusters (facilitating comparison between discrete cell states) and the components of the HPF (facilitating analysis of modular processes shared between cells). For visualization, we embedded the manifold in two dimensions using an optimized tSNE projection from HPF space (e.g. Figure 1). Unless otherwise specified, the following parameters were used: `k=25, k_pooling=10, factorization=“HPF”, n_factors=96, n_genes=2000, mask=(), doublets_action=None, features=“enrichment”`.

Cytograph uses the following open-source packages, and we are grateful to the authors for making such important resources freely available to the community: `numpy, scikit-learn, scipy, networkx, python-louvain, hdbscan, pyyaml, statsmodels, numpy-groupies, tqdm, umap-learn, torch, harmony-pytorch, pynndescent, click, leidenalg, unidip and opentsne.`

#### Distance metric, manifold graph, and pseudo-lineage geodesics

In single-cell analysis, it is typical to construct a nearest-neighbor graph by imposing a distance metric on a reduced-dimensional space of gene expression. This graph is a proxy for the manifold of allowed gene expression states, and can be used for downstream processing such as clustering. However, the standard approach has several issues:

- Nearest neighbors need not be close to each other in any absolute sense. A lone blood cell among neurons will be nearest to some neuron, but not close in any useful sense.
- Distances are not biologically meaningful. For example, “euclidean distance in PCA space” has no intrinsic meaning that can be interpreted as a meaningful distance between cells.
- Neighborhood structure on the manifold graph depends strongly on sampling density

We address these issues in three ways:

- We use Hierarchical Poisson Factorization (HPF) to model the gene expression matrix as a linear combination of factors, and show that this model captures nearly all structure in the expression matrix (Extended Data Figure 3a-d).
- We use the Jensen-Shannon Divergence, also known as the “information radius” as a principled metric that distinguishes “near-identical” cells from “dissimilar cells” (Extended Data Figure 3e-i).
- We define pseudo-lineage trajectories as geodesics along the manifold, which therefore correspond to sets of cells that form a path connected by pairs of cells within a defined maximal information radius (Extended Data Figure 8a-b).

Given a pair of cells *i, j* and their normalized HPF component vectors *θ*_*i*_, *θ*_*j*_ (considered here as probability distributions over the components) the Jensen Shannon Divergence (JSD) is defined as

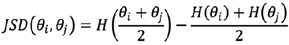

where *H(*·) is the Shannon entropy function. The JSD measures the entropy of the average of two cells, compared to the average entropies of the individual cells. It is a symmetrized version of relative entropy (Kullback-Leibler divergence). For a pair of identical cells, their HPF component values are drawn from identical distributions, the entropy of the average is equal to the average of entropies, and the JSD is zero. For a pair of dissimilar cells, the average of entropies is smaller than the entropy of the average, because information is lost in averaging. A pleasing property of the JSD is the fact that its square root is a true metric (satisfying non-negativity, symmetry, identity of discernibles and triangle inequality), which is required by some clustering and manifold learning algorithms. The JSD can be viewed as defining a small neighborhood around each cell, in which neighbors are near-indistinguishable in an information-theoretic sense. Exploiting this property, we construct a radius nearest neighbor (RNN) graph, by imposing a fixed JSD radius around each cell, and connecting only cells that fall within this radius (for computational efficiency, we cap the maximum number of neighbors to a fixed number, usually 25). This approach addresses several of the deficiencies of the common KNN approach:

- Distances between neighbors have a clear meaning in terms of relative entropy, and informally, cells that are connected in the RNN can be viewed as “near-identical”
- If the manifold is undersampled, the RNN will automatically be sparsely connected, or even disconnected, reflecting missing neighbors
- Regions of the manifold that have been densely sampled will be densely connected
- Connected paths through the RNN are guaranteed to proceed through individual cell neighbors that are near-identical in terms of relative entropy; a more meaningful property than arbitrary distances in PCA space

Next, for inference of longer trajectories along the manifold, we define a pseudo-lineage geodesic as the shortest path between two given cells (e.g. a root cell and a terminally differentiated cell) on the RNN using the JSD as edge weights. The shortest path is therefore a path of small steps, each connecting a pair of near-identical cells. A pseudo-lineage tree of all cells can be constructed by designating a single root cell, and computing the geodesics to all other cells. On this tree, many cells will be internal (branches), some external (leaves) and the root cell will have no predecessor. Cells not connected directly or indirectly to the root cell will not be part of the tree. For this paper, we constructed the pseudo-lineage tree by picking a single cell (#206,003) in the early neuroepithelium arbitrarily as the root.

#### Auto-annotation

After clustering, Cytograph incorporates a step termed auto-annotation that automatically assigns meaningful labels to the clusters (Extended Data Figure 10). Labels are flexible and may denote either specific cell types or general properties such as “cell cycle” or “GABAergic neurotransmission.” Multiple labels can be assigned to the same cluster, and each label can be attributed to multiple clusters. Each label is defined as a text document with a short computer-readable section that consists of four mandatory fields: name, abbreviation, definition and category:

name: `Cranial Neural Crest Cell`

abbreviation: `CNC`

definition: `+Dlx2 +Plp1 +Sox10`

categories: `Ectodermal NeuralCrest Face`

The “abbreviation” and “name” are unique identifiers, whereas “categories” can be used to organize the labels into meta-labels. The “definition” is the most important field and represents the label-assignment rule. It consists of a list of gene names, each prefixed with “+” if the gene must be present or “-” if the gene must be absent. To determine presence or absence, we use Cytograph’s trinarization score^22^. A cluster is assigned a label if and only if the “+” genes are present and the “-” genes are absent. Many different definitions for the same label are possible. To create robust labels that apply to diverse datasets, we favored definitions that comprised short lists of genes that are ideally well-studied. The auto-annotation tags were used for data exploration and to subset the dataset as described below.

Each document additionally contains a free-form section that is used to provide context, justification, and references, which was refined continuously throughout our study and constructed by cross-referencing the literature and our data. As a result, it forms a cumulative body of knowledge about mouse brain cell types and states that can be automatically applied to future datasets, available at https://github.com/linnarsson-lab/auto-annotation-md.

#### Punchcards

Repeating feature selection and dimensionality reduction on a subset of cells often highlights further structure than can be revealed by clustering a large, diverse dataset. Although subsetting the dataset in a supervised way incorporates biological knowledge to expose particularly interesting aspects of the heterogeneity, these decisions quickly become verbose, and the approach requires manual intervention at each level of analysis.

We addressed this issue by using the auto-annotation framework to implement a rule-based splitting procedure. Each level of analysis is defined by a .yaml file termed a punchcard. The name of the punchcard indicates the parent analysis to use as input (e.g. `MidbrainE911.yaml`). Within the file, each subset to analyze is given a name (e.g. `Neurons` and `Glia`) and the clusters to include are determined by the auto-annotation listed in the `include` field. A conditional expression can optionally be specified in the `onlyif` field to further constrain the subset. Punchcards can therefore define a series of biologically meaningful analyses that remain general to the input dataset and analysis parameters. Punchcards can be applied hierarchically to define increasingly fine-grained analyses.

### Post-processing

#### Removal of doublets and contaminated clusters

Doublets were removed by running a modified version of DoubletFinder available in cytograph on each individual sample in the dataset. DoubletFinder scores were then averaged across clusters, and clusters with an average doublet score greater than 0.3 were removed from further analysis. Approximately 20 additional clusters were removed that were manually annotated as blood-contaminated, doublets, or contained fewer than five cells.

#### Merging

A dendrogram was constructed using Cytograph’s `Aggregator` class on the dataset. The dendrogram was cut at a height of 20 to generate groups of similar clusters. For each group, differential expression testing was performed pairwise between the clusters. The package `diffxpy` was used to run t-tests on the log expression values. Genes were considered differentially expressed if 1) *q*-value was less than 0.01, 2) log fold change was greater than 2, and 3) at least 30% of cells expressed the gene in one cluster, and the percentage of expressing cells in the second cluster was less than 70% of the percentage of expressing cells in the first cluster. Clusters with less than one differentially expressed gene were merged. The procedure was repeated until the remaining clusters in each group were distinguishable from one another by at least one differentially expressed gene.

### tSNE of all cells and dendrogram of all clusters

We computed HPF to 30 components and then used the Art of tSNE heuristics^48^ (but with exaggeration=1.5, perplexity=150) to obtain the tSNE layout in Figure 1. The use of a relatively small number of HPF components resulted in a globally smoother and more connected manifold, at the expense of reduced local resolution. This tSNE was used for visualization only; clustering and annotation was performed as described above.

The final dendrogram was generated by running Cytograph’s `Aggregator` class on 942 clusters with parameter mask set to [`‘cellcycle’, ‘sex’, ‘ieg’, ‘mt’`], which excludes cell-cycle, sex-related, immediate-early, and mitochondrial genes from the analysis. Both the cluster dendrogram and the tSNE of all cells were computed with a more recent version of cytograph (commit 8a5518e-81197f5aa9d54565cbce3875738d8dac9).

### Glia analysis

#### Clustering

Cells belonging to clusters 751 and above were pulled from the dataset. The mean expression of each gene was calculated for every cluster, and a new dendrogram was calculated based on the correlation of gene expression between clusters. The resulting dendrogram was cut at a height of 1, resulting in 18 meta-clusters that were subsequently annotated based on gene expression. Independently, a new t-SNE was calculated on this subset of cells using cytograph with the parameters described above except `n_factors`: 50 and mask: [`‘cellcycle’, ‘sex’, ‘ieg’, ‘mt’, ‘ery’`] to mask cell-cycle, sex-related, immediate-early, mitochondrial, and blood-related genes.

#### Differential expression

Differential expression was calculated using the `diffxpy` package. A t-test was calculated on the log gene-expression values normalized by total molecules per cell. Independently, the fraction of positive cells for each gene was calculated for both neurogenic radial glia and glioblasts. Differentially expressed genes were then filtered for those with *q*-values less than 10^−5^ and expressed in at least 10% of either neurogenic radial glia or glioblasts.

#### Cell type scores

Cell-type scores were based on the top 200 enriched genes for astrocytes and OPCs in the adolescent mouse dataset. The score was calculated as the percentage of total genes expressed in each cell that belonged to the enriched set.

#### Region classifier

1 200 cells were sampled from each of six subsets of the dataset: early glia, neurogenic glia, gliogenic glia, OPCs, neuroblasts, and neurons. Equal numbers were sampled from the forebrain, midbrain, and hindbrain. Each of these subsets was used to train a classifier for region that was then tested on the other five groups. The `sklearn GradientBoostingClassifier` was regularized with the parameter `max_features`, and `max_features=50` was chosen after testing the performance of the classifier over a range of values. Default values were used for other parameters. Precision, recall, and F1 scores were calculated separately for forebrain, midbrain, and hindbrain cells and averaged.

### Gastrulation and neural plate analysis

The dataset from Pijuan-Sala et al. was prefiltered by removing cells not classified by the authors. Feature selection using a CV-mean was used to restrict the analysis to the most variable 3500 genes. Analogously 5 223 genes were selected for the E7-8 dataset (2 000 for the primitive streak-stage; non-neural lineage-restricted, and early neuroepithelial). We considered the intersection between these two gene sets. The count tables were imputed by k-nearest neighbor pooling^46^ (`k=30`) and then log-transformed. To place the E7-8 data on the t-SNE embedding for Pijuan-Sala et al., we first computed the similarity (Pearson’s R) between each of the E7-8 cells and the reference cells. Then we assigned to each cell the average of the embedding coordinates computed over the 20 most similar cells in the reference.

The signature score for neuroblast from different regions was obtained considering genes enriched in neuroblasts sampled from forebrain, midbrain and hindbrain. In particular, we considered the neuroblast clusters sampled between E9 and E13 and computed an enrichment score for each of the areas and selected the top 200 genes. Neuroblast clusters were defined using a rule based on the auto-annotations: negative for the cell cycle label and positive for one of the neuronal lineage labels (Glut, GABA, Glyc, etc.). Finally to compute the signature score for each cell we used the same approach adopted by Seurat^49^. We calculate the Z-scores for each of the genes in the set, then average them and normalize the result using a reference computed by drawing random gene samples.

### Spatial localization of organizer-like clusters

We used the R package Voxunt to determine the developing brain regions most likely to host each of the organizer-like cell clusters^40^. The tool constructs a 3D similarity map of the cells using a reference spatial expression atlas that can be interpreted as a putative localization for the cells. Briefly, the tool computes the Spearman correlation between average gene expression of a scRNA-seq cluster and each voxel of a gene expression model constructed from the ISH experiments of the Allen Brain Developmental Atlas^50^. To compute the similarity map we used the same script for E11 and E13, where we call `variable_genes` with `nfeatures=150` and run `voxel_map` and `plot_map` using default parameters.

### In situ sequencing analysis

Channel images were merged into a multichannel tiff image and aligned to a reference round. The channel used for tile alignment was the DAPI stain. Following alignment, the tiles were stitched together using Microscopy Image Stitching Tool (MIST^51^) and subsequently split into non-overlapping 2000×2000 pixel tiles. Each individual tile was then top-hat filtered, the RCPs were segmented and the intensity was measured in each channel. The RCP was then assigned the base with the highest intensity. The public repository for the code used can be found at https://github.com/Moldia/Tools. Gene expression was summarized considering a isotropic grid which step 16 μm and counting the called dots for each gene. The data was smoothed with gaussian filter with a bandwidth of 24 μm. The grid location with a total of called dots smaller than 4 were excluded from the clustering analysis. As a preprocessing before clustering the counts were transformed with the variance stabilizing transform *log*_*2*_*(x+1)*, PCA was computed and the top 40 principal components retained. Finally, we performed a KMeans clustering setting `n_cluster=40`.

### Glioblastoma analysis

To identify aneuploid cells, we calculated the normalized sum of expression on each chromosome, and then fit a 5-component Bayesian Gaussian mixture model to the 24-dimensional chromosome expression data of all cells (Extended Data Figure 9a-c). We then inferred the maximum a posteriori component label for each cell, and the posterior probability. We manually examined these labels for all clusters to determine which components corresponded to aneuploid cells, and which were euploid. We then mapped human to mouse genes and re-clustered and re-annotated the tumor data using cytograph to ensure comparable analyses. To match tumor clusters with reference cell types, we first computed the top 25 enriched genes for each reference cell type (developmental and adolescent). For each reference cluster and each tumor cluster, we then computed a matching score as the product of gene enrichment in the tumor for the 25 top enriched genes in the reference cell type. Thus, a high matching score reflected highly specific expression of the set of genes specifically expressed in the reference cell type. The top-scoring matching cell type was selected for each tumor cluster. For immune cells, we lacked reference data, and therefore called those cell types manually based on well-known markers of macrophages, B cells and T cells.

---

**Extended Data Figure 1.**
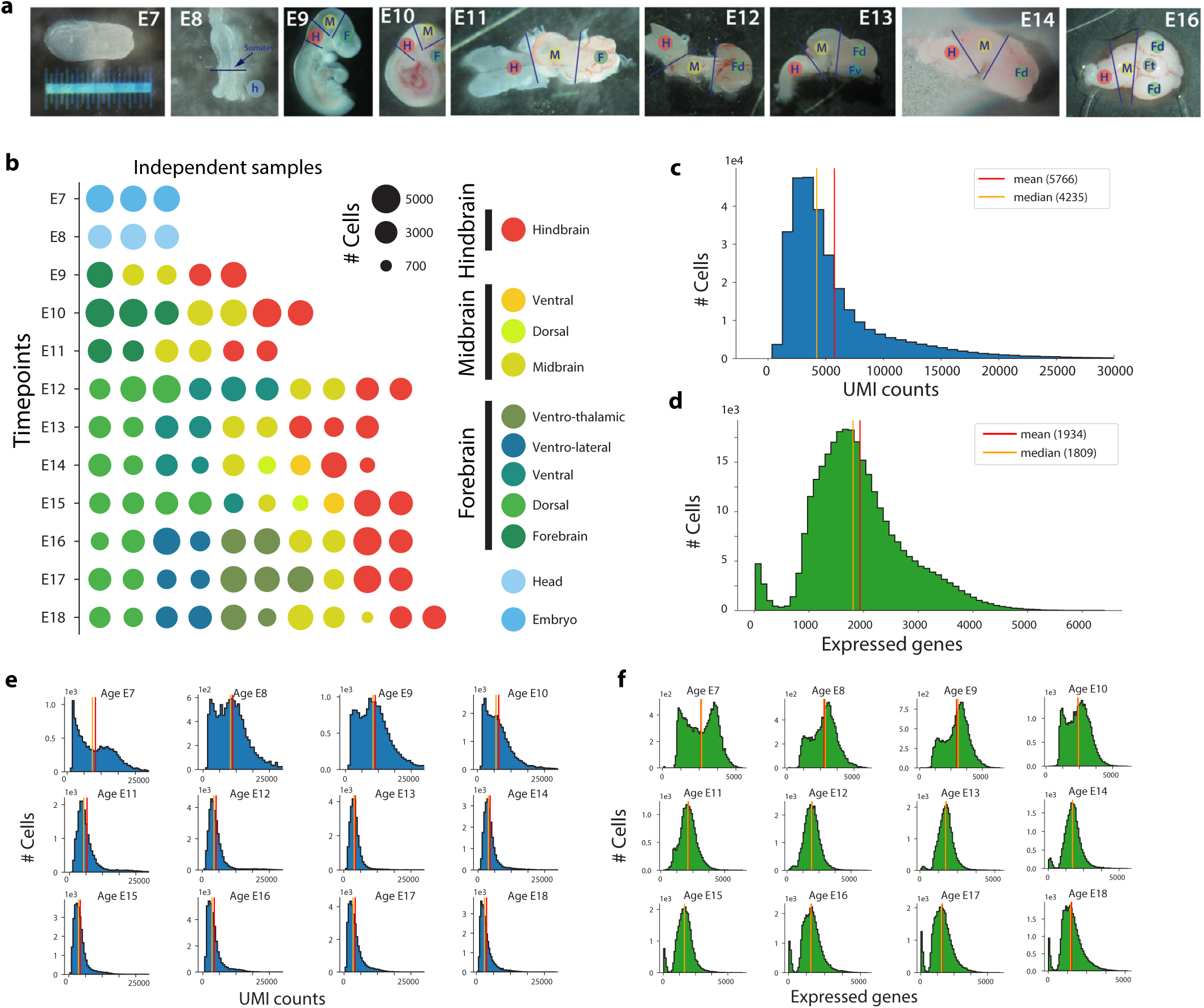
Experimental design and data quality. **a**, Stereo microscope photographs illustrating the tissue dissection strategy. **b**, Samples composing the dataset. The area of each circle is porportional to the number of cells sampled. Color indicates the region dissected. **c**, Histogram showing the distribution of number of UMIs per cell detected accross the entire dataset. **d**, Histogram showing the distribution of number of genes detected per cell accross the entire dataset. **e**, Distribution of UMIs per cell aggregated per age group. **f**, Distribution of number of genes detected per cell aggregated per age group.

**Extended Data Figure 2.**
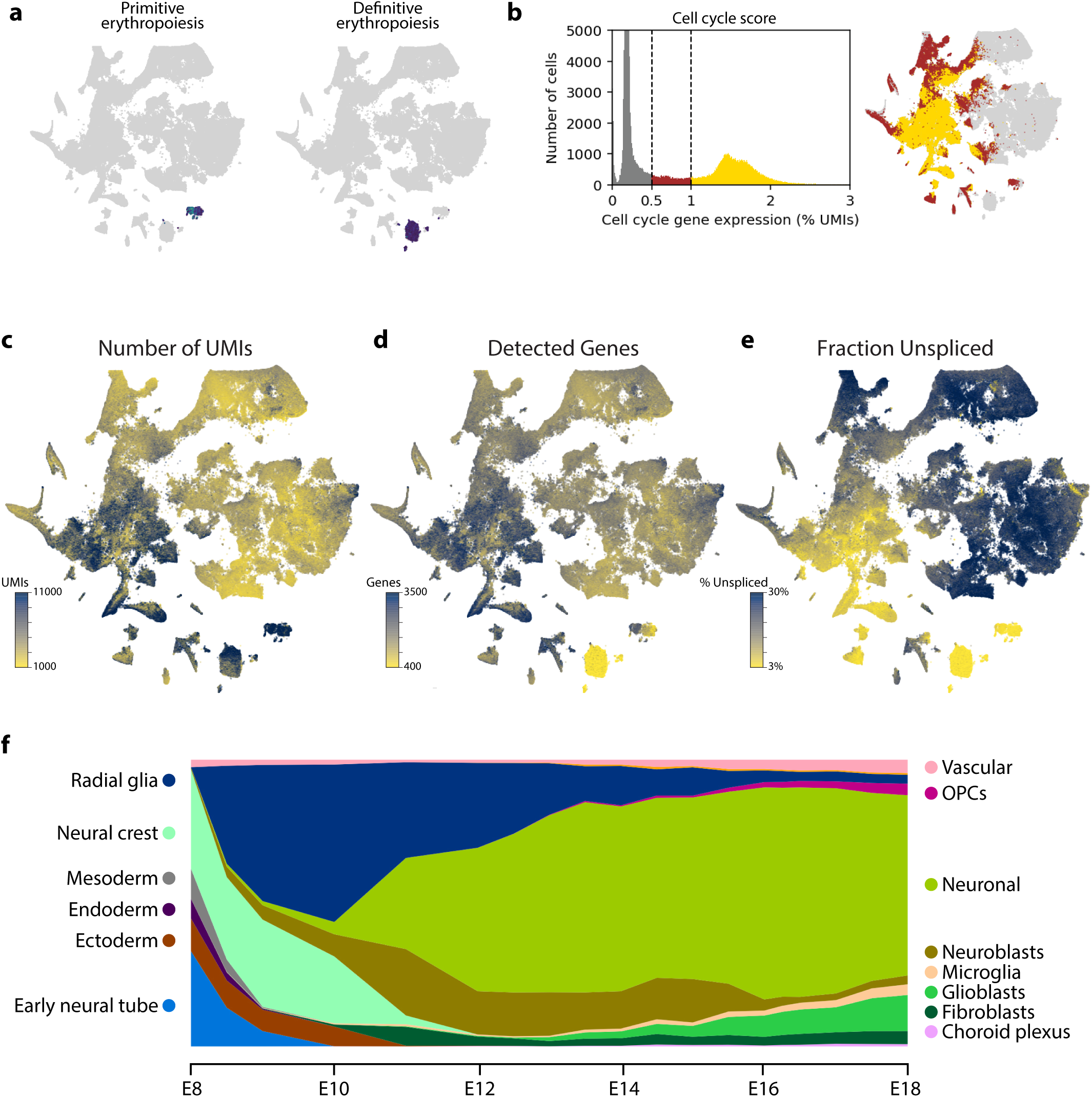
Global properties of gene expression during brain development. **a**, Erythrocyte clusters expressing *Hbb-bh1* (>500 UMIs/cell; primitive erythropoiesis) or *Hbb-bs* (>5000 UMIs/cell; definitive erythropoiesis). **b**, Histogram of cell cycle scores (left) and cell cycle scores indicated on the main tSNE (right). **c**, Distribution of total UMIs per cell. **d**, Distribution of total number of genes detected per cell. **e**, Distribution of the fraction of unspliced reads detected per cell. **f**, Distribution of major classes of cells by gestational age.

**Extended Data Figure 3.**
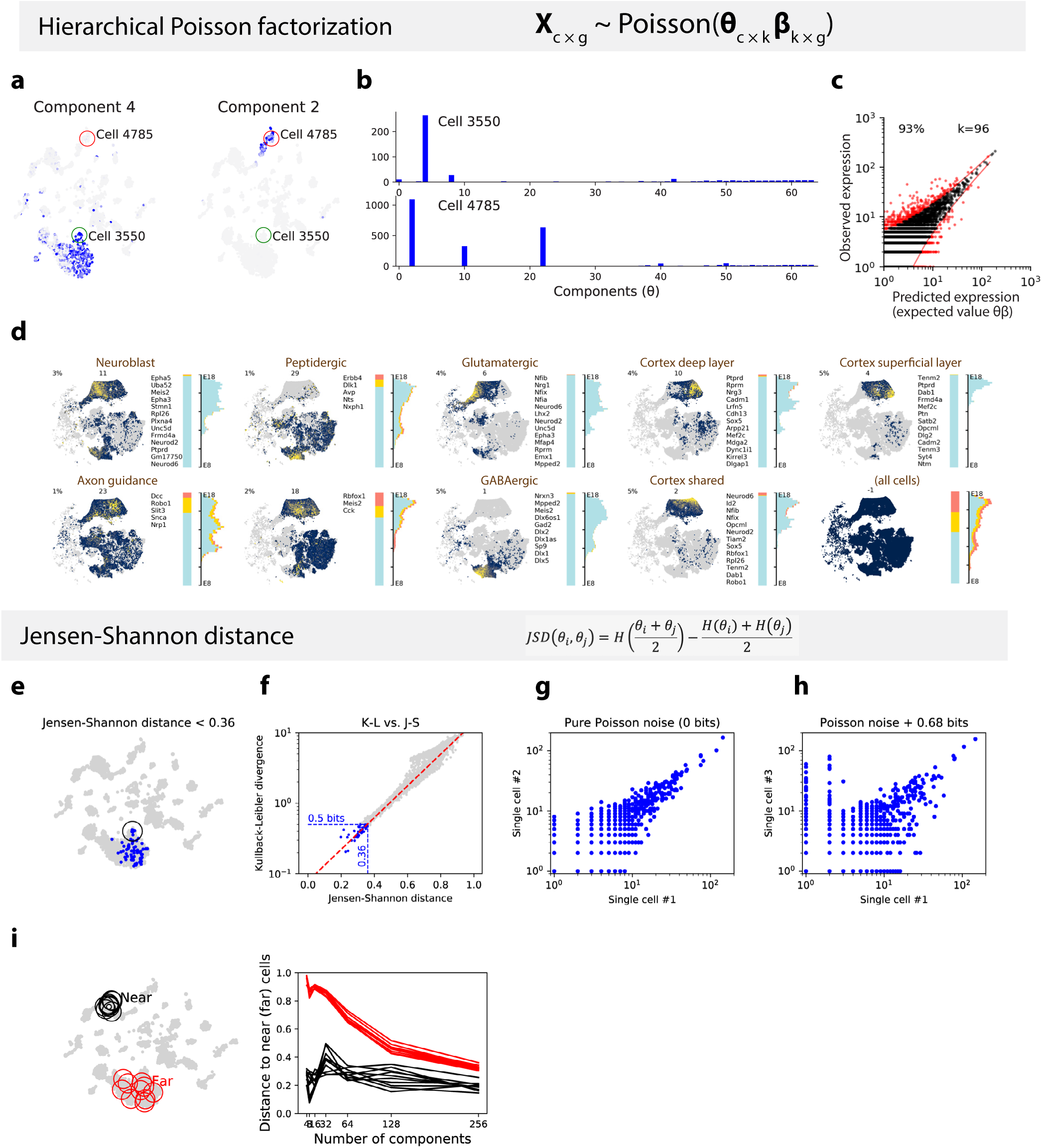
Computational methods. **a**, Illustration of hierarchical Poisson factorization (HPF) on a small subset of cells using 32 components, showing component loadings across cells for components 4 and 2, illustrating the modularity of components. **b**, Loadings of all components for two single cells, illustrating the sparseness of components; each cell is approximated by a small number of non-negative components. **c**, Scatterplot showing observed expression of all genes in all cells, versus predicted expression based on 96 HPF components. 93% of datapoints were located inside the 95% confidence intervals of the Poisson distribution, demonstrating the accuracy of the HPF representation. **d**, HPF analysis of all neuronal cells from the full dataset using 32 components, showing that HPF captures essential features of forebrain neurogenesis. Each subplot shows a tSNE colored by component loading, a list of genes with high loading in the component, and bar chart and histogram showing the regional distributions of cells (cf. the distribution of all neuronal cells, lower right). Nine selected components are shown, related to forebrain based on the regional distribution of cells with high loadings. **e**, Example of cells (blue) that fall within a Jensen-Shannon distance of 0.36 of a single cell (black circle), corresponding to 0.5 bits of Shannon entropy. **f**, The relationship between the Kullback-Leibler divergence and the Jensen-Shannon distance, showing that in this case 0.36 JSD corresponds to 0.5 bits. **g**, Two virtual cells — Poisson samples drawn from a single HPF vector — showing the distribution of pure Poisson expression noise. **h**, The effect of switching two HPF components, resulting in a change in expression of many genes and a Jensen-Shannon distance of 0.68 bits. **i**, The curse of dimensionality. As the number of HPF components is increased, the difference in Jensen-Shannon distance between nearest-neighbors and far-away cells on the manifold decreases.

**Extended Data Figure 4.**
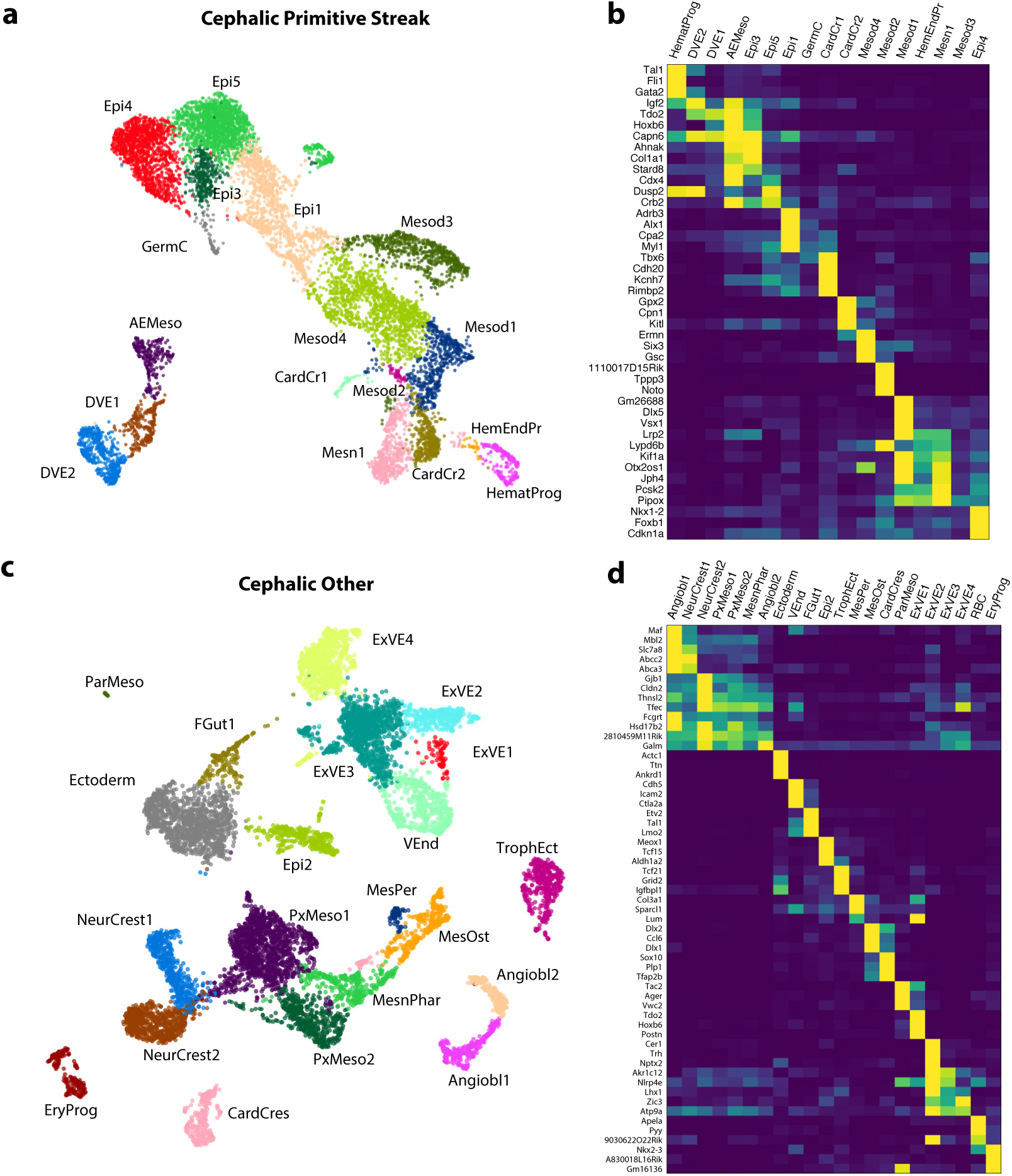
Neurulation stage. **a**, tSNE embedding of cells related to the cephalic primitive streak, colored and labelled by cluster identity. **b**, Heatmap showing genes enriched in the clusters shown in a. **c**, tSNE embedding of cells related to other lineages, colored and labelled by cluster identity. **d**, Heatmap showing genes enriched in the clusters shown in (c).

**Extended Data Figure 5.**
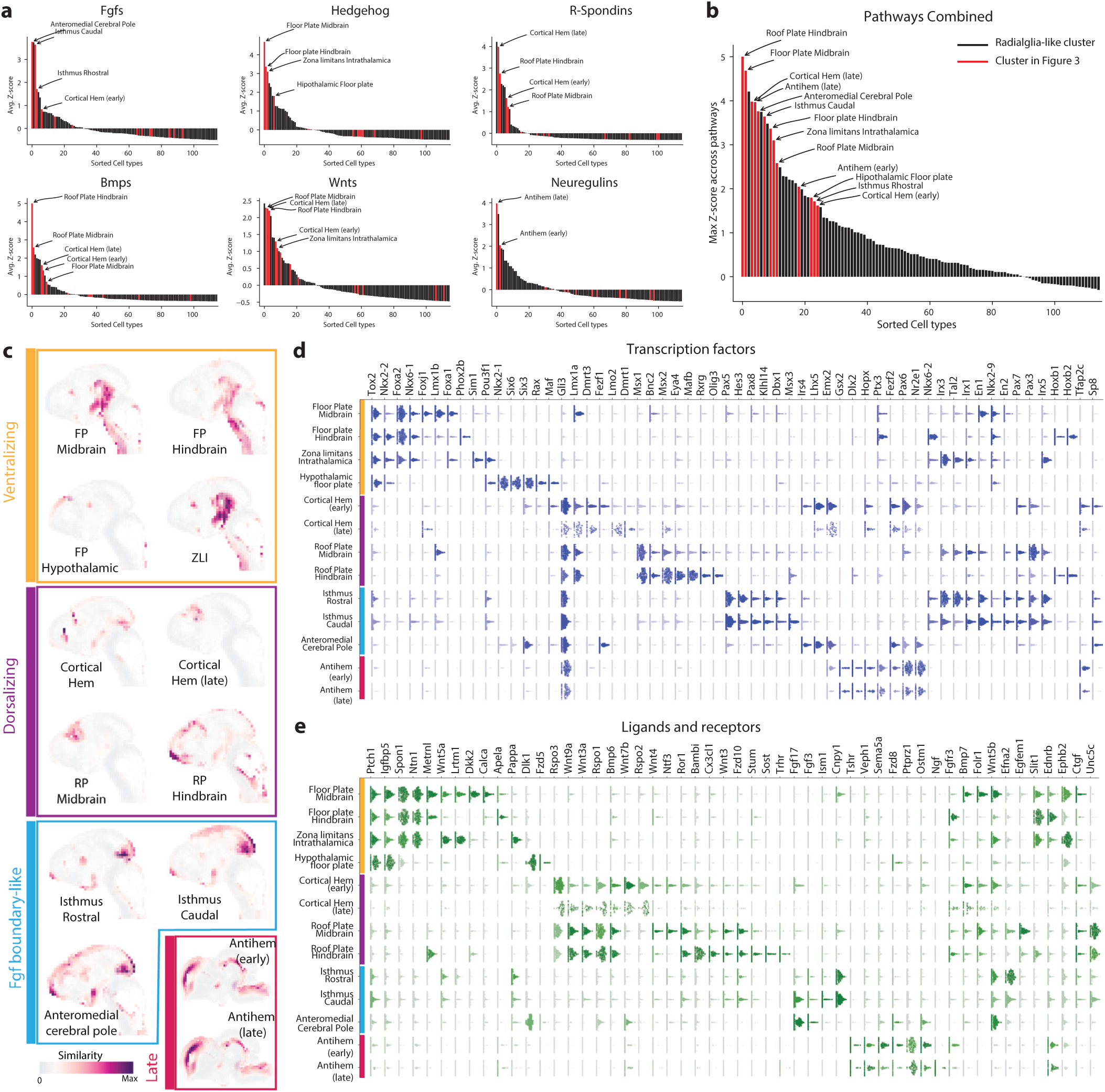
Secondary organizers. **a**, Cell types sorted on the basis the average Z-score of morphogens genes grouped in categories on the basis of gene family and the neurulation literature. Hedgehog (*Shh* and *Ptch1*), R-spondins (*Rspo1, Rspo2, Rspo3* and *Rspo4*), Wnts (*Wnt1, Wnt3a, Wnt5a* and *Wnt8b*), Neuregulins (*Nrg1, Nrg3* and *Fgf7*), Fgfs (*Fgf8, Fgf15, Fgf17* and *Fgf18*), Bmps (*Bmp6, Bmp7* and *Gdf7*). Cell types displayed in red were localized and analyzed in Figure 3 and in the panels below. **b**, Summary of the plots in (a) showing the maximum average Z-score achieved. **c**, Putative localization of the cell populations by gene expression matching with the Allen Brain ISH Atlas. Images show a 2D projection of a 3D voxel map colored by the Spearman correlation coefficient between the voxel expression and the cluster. **d**, Beeswarm plots showing the expression of a selection of transcription factors that were found enriched in the different organizer cell populations. **e**, Beeswarm plots showing the gene expression of enriched receptors and ligands whose distribution in relation to organizers had not been described. Genes in (d) and (e) were selected for HybISS profiling.

**Extended Data Figure 6.**
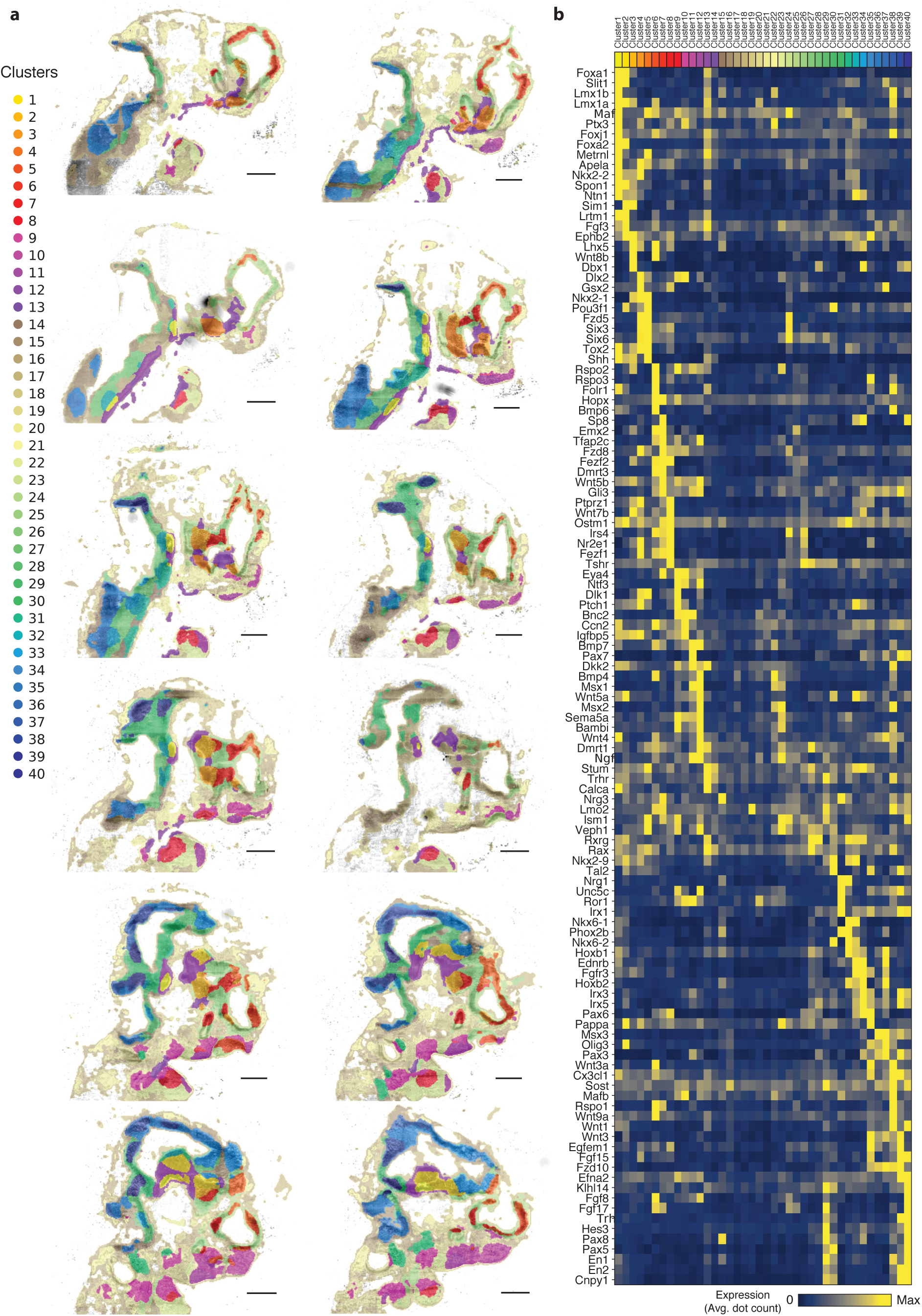
In situ sequencing analysis. **a**, Twelve sagittal sections of the E10.5 embryo with pixels colored by the clusters defined using the 119 genes measured by HybSS. Cluster colors are overlaid onto the DAPI channel plotted in grayscale. The sections span the neural tube from lateral (above) to medial position (below). Scale bars, 480 μm. **b**, Heatmap displaying the gene expression of each of the 119 genes measured by HybSS in the embryonic regions defined by clustering. Expression was quantified counting dots on a grid with 16 ☒m steps and the dot count was averaged across all the grid points assigned to each cluster.

**Extended Data Figure 7.**
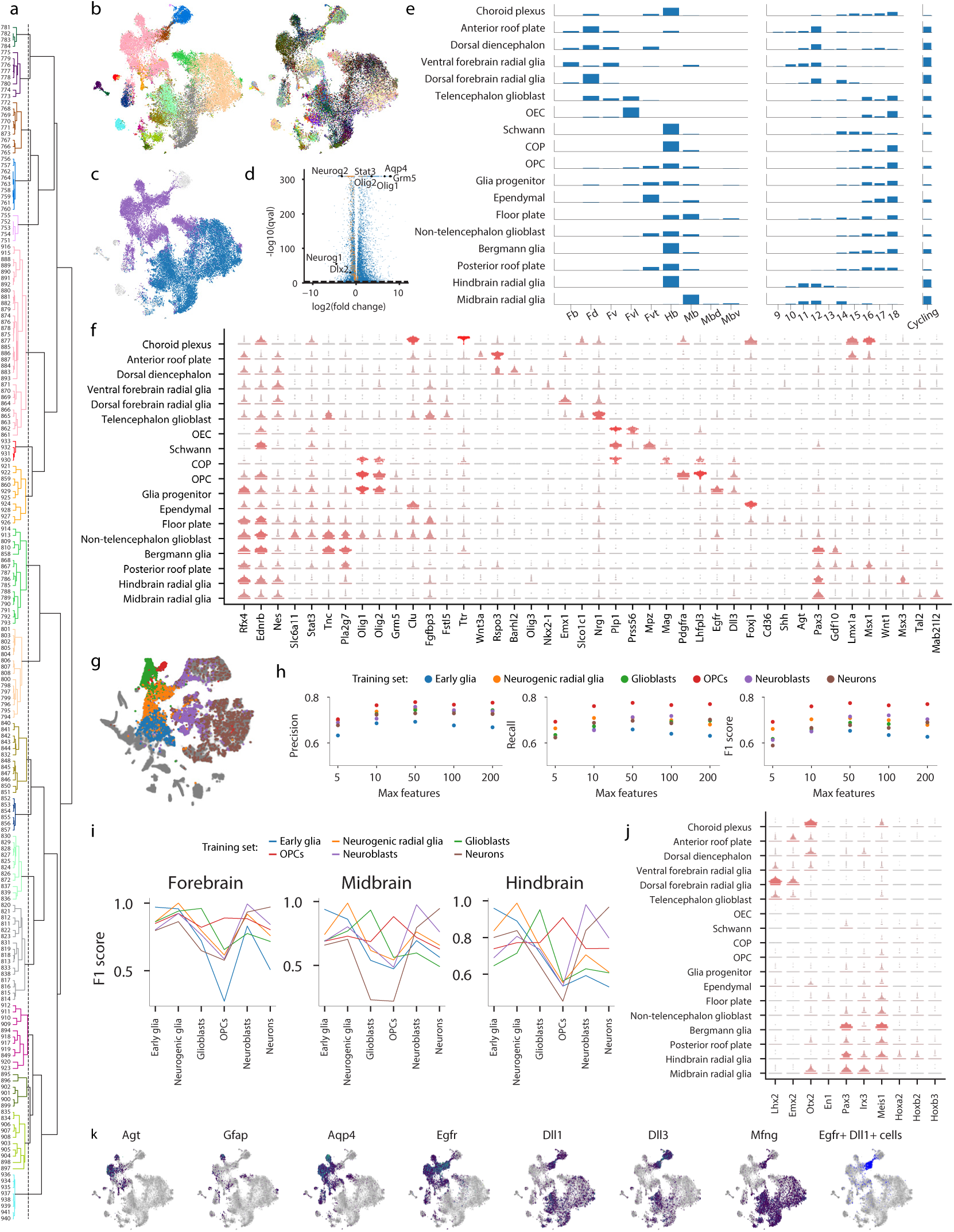
Glial diversity. **a**, Cells from clusters 751 and above were pulled from the complete dataset. Data were aggregated by cluster, features were selected, and a dendrogram was built using correlation distance. The dendrogram was then cut at a height of 1 (dotted line). Colors indicate the 18 resultant meta-clusters. **b**, A t-SNE embedding was calculated on the same subset of cells. Cells are colored by their original cluster (left) and meta-cluster cut from the dendrogram (right). **c**, Cells are colored by their labels for differential gene-expression testing: purple-glioblast; blue-neurogenic; grey - not included in testing. **d**, Select genes are annotated on a volcano plot illustrating differential expression between neurogenic and gliogenic radial glia. The dotted line denotes the significance threshold (*q* = 10^−5^). The range of the x-axis was chosen to capture statistically significant genes. Cell-cycle genes are colored orange. **e**, For each meta-cluster the height of the bar indicates the percentage of cells from each tissue (left) or each embryonic stage (middle) or that are cycling (right). Abbreviations are Fb - forebrain, Fd - dorsal forebrain, Fv - ventral forebrain, Fvl - ventrolateral forebrain, Fvt - ven-trothalamic forebrain, Hb - hindbrain, Mb - midbrain, Mbd - dorsal midbrain, Mbv - ventral midbrain. **f**, Expression dot plots are shown for select genes. **g**, Cells selected for region classification are colored: blue - early glia, orange - neurogenic glia, green - glioblasts, red - OPCs, purple - neuroblasts, brown - neurons. Grey cells were not selected. **h**, For a range of parameter values (max_features), a gradient boosting classifier was fit to each training set and tested on the five remaining training sets. Average precision, recall, and F1 scores are plotted for each parameter value and training set. **i**, F1 scores for each gradient boosting classifier (max_features = 50) are shown for forebrain, midbrain, and hindbrain cells. The color of each line indicates which cells were used to train the classifier (colored as in g). **j**, Feature importances for each gradient boosting classifier were used to rank genes, and the intersection of the 100 most important genes for all six classifiers is shown on the x-axis. Expression dot plots are shown for each of these genes. **k**, Cells are colored by log gene expression for the indicated genes. Grey indicates no expression. Cells that are positive for both *Egfr* and *Dll1* are colored blue.

**Extended Data Figure 8.**
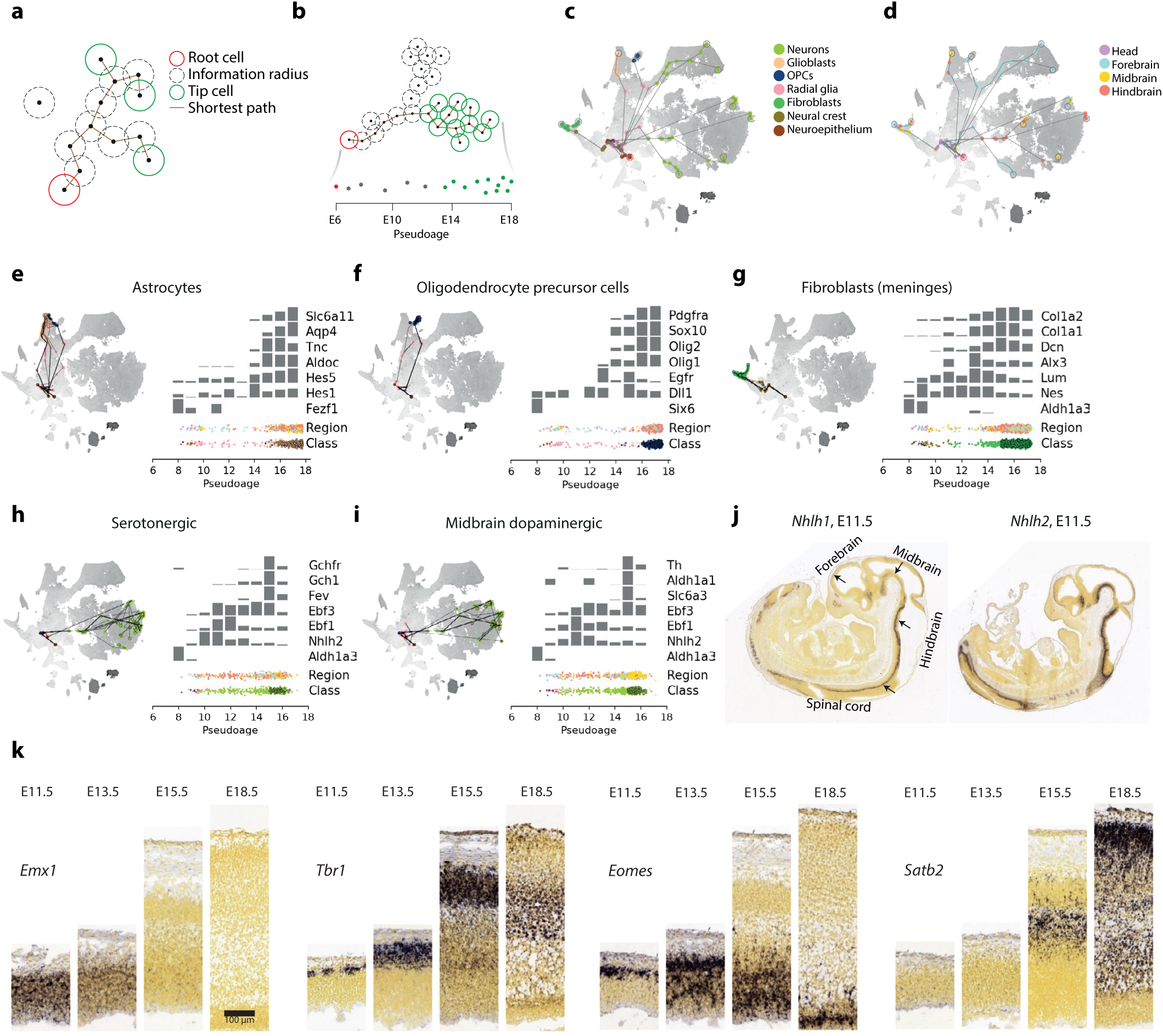
Pseudolineage analysis and neurogenesis. **a**, Pseudolineage tree algorithm, computing shortest paths to the root cell passing only through neighbors inside the information radius. **b**, Isolating the pseudolineages terminating in a selected cluster, and projecting to the pseudoage axis. **c**, Ten selected pseudolineages colored by major class, on the tSNE of Figure 1 with greyscale showing the geodesic distance to the root from every cell. **d**, The same ten selected pseudolineages colored by tissue. **e-i**, Pseudolin-eages of astrocytes (e), OPCs (f), fibroblasts (g), hindbrain serotonergic neurons (h), and midbrain dopaminergic neurons (i). Each plot shows a randomly selected subset of pseduolineages terminating in the indicated clusters, as well as expression of selected genes in pseudoage bins along the lineage (calculated for all cells in the lineage). The region and class of each cell is indicated at the bottom. Isolation of pseudolineages terminating in a selected cluster, and projection to the pseudoage axis. **j**, Astrocyte and OPC scores. **k**, RNA in situ hybridization of mouse embryonic brain at the indicated timepoints, showing genes relevant to the cortical lineage (image credit: Allen Institute). Each subpanel shows a strip from ventricular zone to pia at four different ages.

**Extended Data Figure 9.**
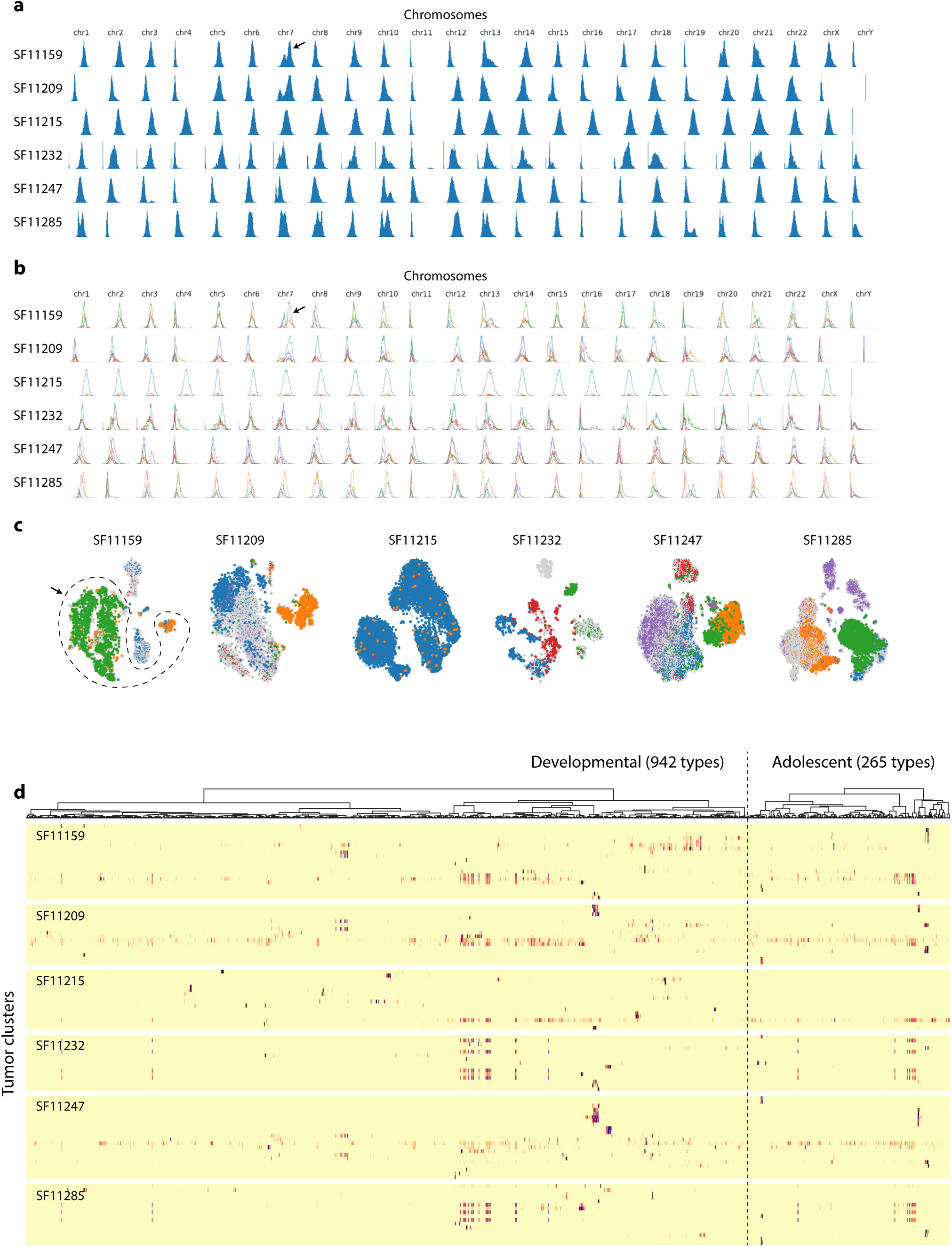
Glioblastoma. **a**, Histograms of total gene expression per cell per chromosome, for each tumor sample. Arrow indicates an amplification of chromosome 7, the most common aneuploidy in glioblastoma. **b**, Fitting a gaussian mixture model to the 24-dimensional chromosome expression data for each cell, with five components. Arrow indicates two components (green and orange) that fit the amplification of chromosome 7. **c**, tSNE plots of each of the tumor samples, colored by the gaussian mixture model prediction for each cell. The size of the cell indicates the probability of the prediction. Arrow and dashed outline indicates cells that likely carry the amplification of chromosome 7 in sample SF11159. **d**, Heatmaps showing the matching score between reference cell types (columns, ordered as in Figure 1 for developmental types, and as in Figure 1 of Zeisel et al. for adolescent types) and tumor clusters (rows). For each tumor cluster (row), the identity of the best-matching cluster is shown in Figure 5.

**Extended Data Figure 10.**
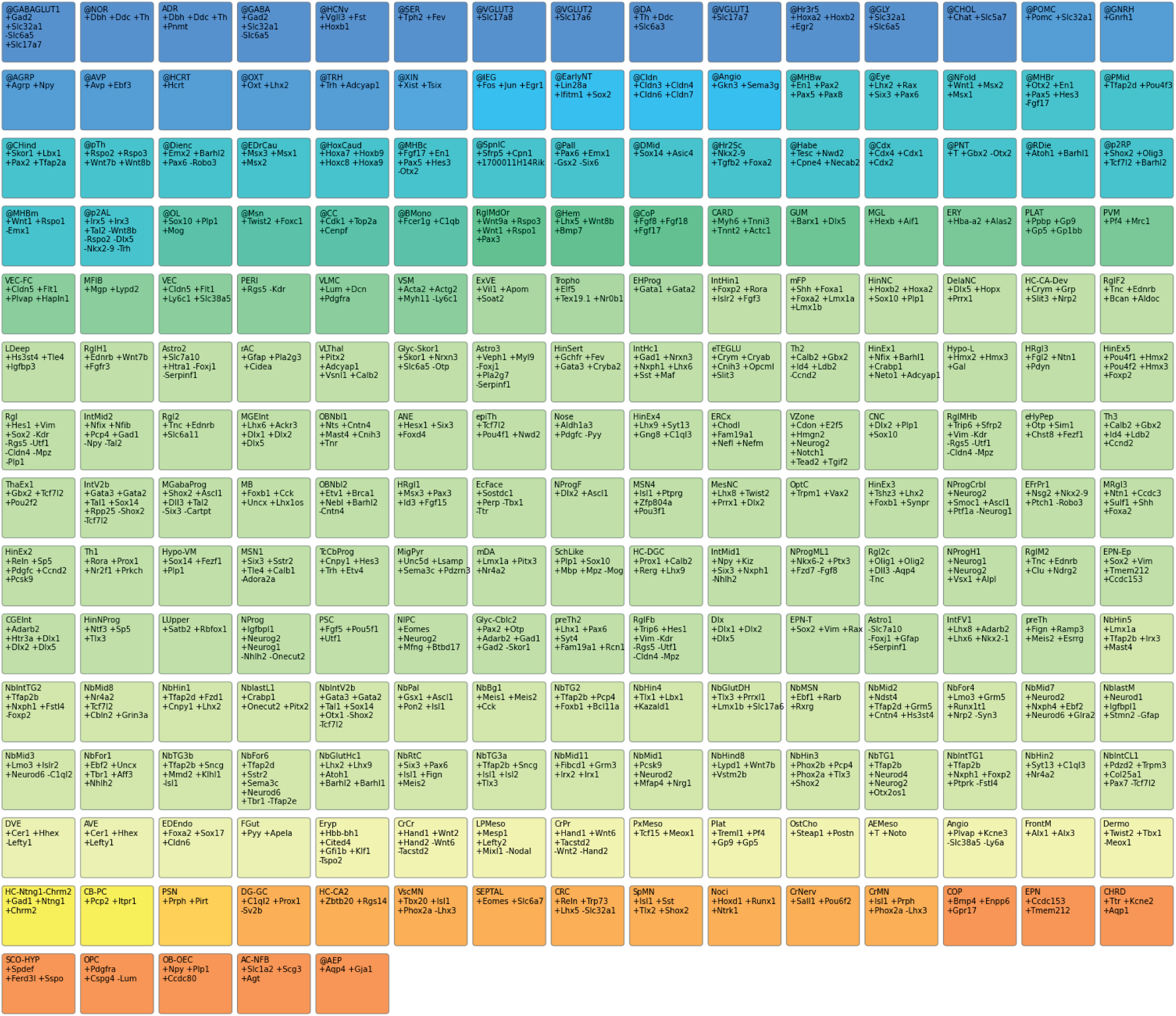
Auto-annotation labels. Each patch shows an auto-annotation label and its definition. (+) indicates a gene required to be expressed, and (-) indicates a gene required to not be expressed for the label to apply. Colors indicate broad categories of labels. All 215 auto-annotation labels used in this paper are shown, and detailed documentation for each label is available at https://github.com/linnarsson-lab/auto-annotation-md/.

## References

1. Sadler, T. W. Embryology of neural tube development. Am. J. Med. Genet. Part C Semin. Med. Genet. 135C, 2–8 (2005).

2. Camp, J. G. et al. Human cerebral organoids recapitulate gene expression programs of fetal neocortex development. Proc. Natl. Acad. Sci. 112, 15672–15677 (2015).

3. Pollen, A. A. et al. Molecular Identity of Human Outer Radial Glia during Cortical Development. Cell 163, 55–67 (2015).

4. Johnson, M. B. & Walsh, C. A. Cerebral cortical neuron diversity and development at single-cell resolution. Curr. Opin. Neurobiol. 42, 9–16 (2017).

5. Nowakowski, T. J. et al. Spatiotemporal gene expression trajectories reveal developmental hierarchies of the human cortex. Science (80-.). 358, 1318–1323 (2017).

6. Zhong, S. et al. A single-cell RNA-seq survey of the developmental landscape of the human prefrontal cortex. Nature 555, 524–528 (2018).

7. Polioudakis, D. et al. A Single-Cell Transcriptomic Atlas of Human Neocortical Development during Mid-gestation. Neuron 103, 785–801.e8 (2019).

8. Yuzwa, S. A. et al. Developmental Emergence of Adult Neural Stem Cells as Revealed by Single-Cell Transcriptional Profiling. Cell Rep. 21, 3970–3986 (2017).

9. Hochgerner, H., Zeisel, A., Lönnerberg, P. & Linnarsson, S. Conserved properties of dentate gyrus neurogenesis across postnatal development revealed by single-cell RNA sequencing. Nat. Neurosci. 21, 290–299 (2018).

10. Cao, J. et al. The single-cell transcriptional landscape of mammalian organogenesis. Nature 566, 496–502 (2019).

11. Telley, L. et al. Temporal patterning of apical progenitors and their daughter neurons in the developing neocortex. Science (80-.). 364, eaav2522 (2019).

12. Habib, N. et al. Div-Seq: Single-nucleus RNA-Seq reveals dynamics of rare adult newborn neurons. Science (80-.). 353, 925–928 (2016).

13. Zhong, S. et al. Decoding the development of the human hippocampus. Nature 577, 531–536 (2020).

14. Kee, N. et al. Single-Cell Analysis Reveals a Close Relationship between Differentiating Dopamine and Subthalamic Nucleus Neuronal Lineages. Cell Stem Cell 20, 29–40 (2017).

15. La Manno, G. et al. Molecular Diversity of Midbrain Development in Mouse, Human, and Stem Cells. Cell 167, 566–580.e19 (2016).

16. Tiklová, K. et al. Single-cell RNA sequencing reveals midbrain dopamine neuron diversity emerging during mouse brain development. Nat. Commun. 10, 581 (2019).

17. Rosenberg, A. B. et al. Single-cell profiling of the developing mouse brain and spinal cord with split-pool barcoding. Science (80-.). 360, 176–182 (2018).

18. Carter, R. A. et al. A Single-Cell Transcriptional Atlas of the Developing Murine Cerebellum. Curr. Biol. 28, 2910–2920.e2 (2018).

19. Huisman, C. et al. Single cell transcriptome analysis of developing arcuate nucleus neurons uncovers their key developmental regulators. Nat. Commun. 10, 3696 (2019).

20. Guo, Q. & Li, J. Y. H. Defining developmental diversification of diencephalon neurons through single cell gene expression profiling. Development 146, dev174284 (2019).

21. Zywitza, V., Misios, A., Bunatyan, L., Willnow, T. E. & Rajewsky, N. Single-Cell Transcriptomics Characterizes Cell Types in the Subventricular Zone and Uncovers Molecular Defects Impairing Adult Neurogenesis. Cell Rep. 25, 2457–2469.e8 (2018).

22. Zeisel, A. et al. Molecular Architecture of the Mouse Nervous System. Cell 174, 999–1014.e22 (2018).

23. Marques, S. et al. Transcriptional Convergence of Oligodendrocyte Lineage Progenitors during Development. Dev. Cell 46, 504–517.e7 (2018).

24. Weng, Q. et al. Single-Cell Transcriptomics Uncovers Glial Progenitor Diversity and Cell Fate Determinants during Development and Gliomagenesis. Cell Stem Cell 24, 707–723.e8 (2019).

25. Marques, S. et al. Oligodendrocyte heterogeneity in the mouse juvenile and adult central nervous system. Science (80-.). 352, (2016).

26. Vanlandewijck, M. et al. A molecular atlas of cell types and zonation in the brain vasculature. Nature 554, 475–480 (2018).

27. Siegenthaler, J. A. & Pleasure, S. J. Meninges and Vasculature. in Patterning and Cell Type Specification in the Developing CNS and PNS 835–849 (Elsevier, 2013). doi: 10.1016/B978-0-12-397265-1.00087-3

28. Niwa, H., Miyazaki, J. & Smith, A. G. Quantitative expression of Oct-3/4 defines differentiation, dedifferentiation or self-renewal of ES cells. Nat. Genet. 24, 372–376 (2000).

29. Burtscher, I. & Lickert, H. Foxa2 regulates polarity and epithelialization in the endoderm germ layer of the mouse embryo. Development 136, 1029–1038 (2009).

30. Li, L. et al. Location of transient ectodermal progenitor potential in mouse development. Dev. 140, 4533–4543 (2013).

31. Zhang, X. et al. Pax6 Is a Human Neuroectoderm Cell Fate Determinant. Cell Stem Cell 7, 90–100 (2010).

32. Hatakeyama, J. Hes genes regulate size, shape and histogenesis of the nervous system by control of the timing of neural stem cell differentiation. Development 131, 5539–5550 (2004).

33. Pijuan-Sala, B. et al. A single-cell molecular map of mouse gastrulation and early organogenesis. Nature (2019). doi: 10.1038/s41586-019-0933-9

34. Tamai, H. et al. Pax6 transcription factor is required for the interkinetic nuclear movement of neuroepithelial cells. Genes to Cells 12, 983–996 (2007).

35. Javali, A. et al. Co-expression of Tbx6 and Sox2 identifies a novel transient neuromesoderm progenitor cell state. Dev. 144, 4522–4529 (2017).

36. Tzouanacou, E., Wegener, A., Wymeersch, F. J., Wilson, V. & Nicolas, J.-F. Redefining the Progression of Lineage Segregations during Mammalian Embryogenesis by Clonal Analysis. Dev. Cell 17, 365–376 (2009).

37. Murdoch, J. N., Eddleston, J., Leblond-Bourget, N., Stanier, P. & Copp, A. J. Sequence and expression analysis of Nhlh1: A basic helix-loop-helix gene implicated in neurogenesis. Dev. Genet. 24, 165–177 (1999).

38. Alves dos Santos, M. T. & Smidt, M. P. En1 and Wnt signaling in midbrain dopaminergic neuronal development. Neural Dev. 6, 23 (2011).

39. Schwarz, M. et al. Pax2/5 and Pax6 subdivide the early neural tube into three domains. Mech. Dev. 82, 29–39 (1999).

40. Fleck, J. S., He, Z., Boyle, M. J., Camp, J. G. & Treutlein, B. Resolving brain organoid heterogeneity by mapping single cell genomic data to a spatial reference. 2507, 1–9 (2020).

41. Gyllborg, D. et al. Hybridization-based In Situ Sequencing (HybISS): spatial transcriptomic detection in human and mouse brain tissue. bioRxiv 2020.02.03.931618 (2020). doi: 10.1101/2020.02.03.931618

42. Miller, F. D. & Gauthier, A. S. Timing Is Everything: Making Neurons versus Glia in the Developing Cortex. Neuron 54, 357–369 (2007).

43. Couturier, C. P. et al. Single-cell RNA-seq reveals that glioblastoma recapitulates normal brain development. bioRxiv 449439 (2018). doi: 10.1101/449439

44. Neftel, C. et al. An Integrative Model of Cellular States, Plasticity, and Genetics for Glioblastoma. Cell 178, 835–849.e21 (2019).

45. Bhaduri, A. et al. Outer Radial Glia-like Cancer Stem Cells Contribute to Heterogeneity of Glioblastoma. Cell Stem Cell 26, 48–63.e6 (2020).

46. La Manno, G. et al. RNA velocity of single cells. Nature 560, 494–498 (2018).

47. Gopalan, P., Hofman, J. M. & Blei, D. M. Scalable Recommendation with Hierarchical Poisson Factorization. in UAI 326–335 (2015).

48. Kobak, D. & Berens, P. The art of using t-SNE for single-cell transcriptomics. Nat. Commun. 10, (2019).

49. Satija, R., Farrell, J. a, Gennert, D., Schier, A. F. & Regev, A. Spatial reconstruction of single-cell gene expression data. Nat. Biotechnol. (2015). doi: 10.1038/nbt.3192

50. Lein, E. S. et al. Genome-wide atlas of gene expression in the adult mouse brain. Nature 445, 168–176 (2007).

51. Chalfoun, J. et al. MIST: Accurate and Scalable Microscopy Image Stitching Tool with Stage Modeling and Error Minimization. Sci. Rep. 7, 1–10 (2017).

